# Glial-dependent clustering of voltage-gated ion channels in Drosophila precedes myelin formation

**DOI:** 10.1101/2023.01.09.523229

**Authors:** Simone Rey, Henrike Ohm, Frederieke Moschref, Dagmar Zeuschner, Christian Klämbt

## Abstract

Neuronal information conductance depends on transmission of action potentials. The conductance of action potentials is based on three physical parameters: The axial resistance of the axon, the axonal insulation by glial membranes, and the positioning of voltage-gated ion channels. In vertebrates, myelin and channel clustering allow fast saltatory conductance. Here we show that in *Drosophila melanogaster* voltage-gated sodium and potassium channels, Para and Shal, co-localize and cluster in an area of motor axons resembling the axon initial segment. Para but not Shal localization depends on peripheral glia. In larvae, relatively low levels of Para channels are needed to allow proper signal transduction and nerves are simply wrapped by glial cells. In adults, the concentration of Para at the axon initial segment increases. Concomitantly, these axon domains are covered by a mesh of glial processes forming a lacunar structure that serves as an ion reservoir. Directly flanking the voltage-gated ion channel rich axon segment, the lacunar structures collapse forming a myelin-like insulation. Thus, Drosophila development may reflect the evolution of myelin which forms in response to increased levels of clustered voltage-gated ion channels.

**One-Sentence Summary:** Evolution of saltatory conductance is mirrored in fly development where glia dependent clustering of voltage-gated ion channels precedes myelination.

## Introduction

A functional nervous system requires the processing and transmission of information in the form of changing membrane potentials. To convey information along axons, neurons generate action potentials by opening of evolutionarily conserved voltage-gated sodium and potassium channels (Moran et al., 2015). Once an action potential is generated, it travels towards the synapse and the speed of information transfer is of obvious importance. It is long established that axonal conductance velocity depends on the resistance within the axon, which inversely correlates with its diameter, and the resistance across the axonal membrane, which is increased by extensive glial wrapping. Furthermore, spacing of voltage-gated ion channels contributes to axonal conduction velocity (Eshed-Eisenbach and Peles, 2019; Freeman et al., 2016; Hodgkin and Huxley, 1952). In vertebrates, unmyelinated axons generally show a small diameter with evenly distributed voltage-gated ion channels along their plasma membrane, and in consequence their conductance velocity is slow (Castelfranco and Hartline, 2015). To speed up conductance, axons grow to a larger diameter and show a clustering of voltage-gated ion channels at the axon initial segment and the nodes of Ranvier which together with the insulating glial-derived myelin sheet allows fast saltatory conductance (Arancibia-Cárcamo et al., 2017; Castelfranco and Hartline, 2015; Cohen et al., 2019; Dutta et al., 2018; Eshed-Eisenbach and Peles, 2019).

In invertebrates, mechanisms to increase conductance speed are thought to be limited by radial axonal growth, as seen in the giant fiber system of Drosophila or the giant axon of the squid (Allen et al., 2006; Hartline and Colman, 2007). No saltatory conductance has been described for invertebrates and it is assumed that voltage-gated ion channels distribute relatively evenly along axonal membranes. Nevertheless, myelin-like structures were found in several invertebrate species, including annelids, crustacean and insects (Coggeshall and Fawcett, 1964; Davis et al., 1999; Günther, 1976; Hama, 1959; 1966; Hess, 1958; Heuser and Doggenweiler, 1966; Levi et al., 1966; Roots, 2008; Roots and Lane, 1983; Wigglesworth, 1959; Wilson and Hartline, 2011a; b). However, it is unknown whether such myelin-like structures also impact the distribution of ion channels.

To address how glial cells affect axonal conductance velocity we turned to Drosophila. In the larvae, peripheral axons are engulfed by a single glial wrap resembling Remak fibers in the mammalian PNS (Matzat et al., 2015; Nave and Werner, 2014; Stork et al., 2008). In addition to insulating axons, we found that glial cells promote radial axonal growth which show a severe reduction in conductance velocity stronger than predicted by the Hodgkin and Huxley equation (Hodgkin and Huxley, 1952; Kottmeier et al., 2020). Thus, wrapping glia might control axonal localization of voltage-gated ion channels.

## Results

### Distribution of the voltage gated sodium channel Para

The Drosophila genome harbors only one voltage-gated sodium channel called Paralytic (Para), which is required for the generation of all action potentials (Kroll et al., 2015). To study the localization of Para and to test whether Drosophila glia affects its localization we and others tagged the endogenous *para* locus with all predicted isoforms being modified (Ravenscroft et al., 2020; Venken et al., 2011) (Figure 1A). In *para*^*mCherry*^ flies, monomeric Cherry (mCherry) is added at the Para N-terminus (Figure 1A). Homozygous or hemizygous *para*^*mCherry*^ flies are viable with only mildly affected channel function (Figure 1B). Para^mCherry^ localizes along many CNS and PNS axons of the larval and the adult nervous system (Figure 1C,D; see Figure S1A-C).

**Figure 1.**
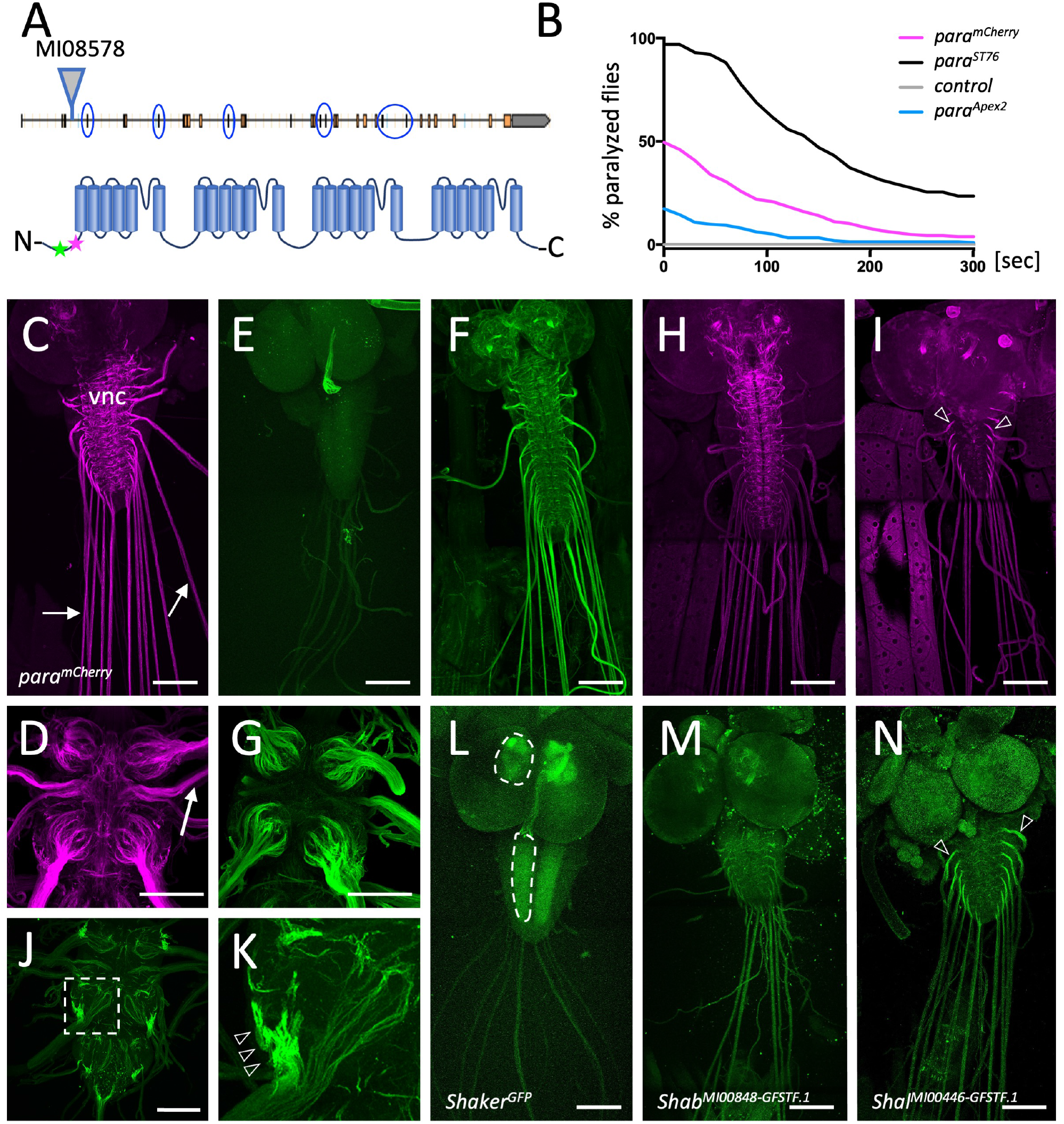
Differential localization of voltage-gated ion channels in the Drosophila nervous system. **(A)** Schematic view on the *para* gene. Alternative splicing at the circled exons results in the generation of more than 60 Para isoforms. All isoforms share a common N-terminus. Here, the MiMIC insertion *MI08578* allows tagging of the endogenous *para* gene. The peptide sequence AEHEKQKELERKRAEGE (position 33-49) that was used for immunization is indicated by a green star, the MiMIC insertion at position 50 is indicated by a magenta star. **(B)** Homozygous *para*^*mCherry*^, *para*^*Apex2*^ or *para*^*ST76*^ flies were tested for temperature induced paralysis. The recovery time is indicated. **(C)** Third instar larval *para*^*mCherry*^ nervous system stained for Cherry localization. Para^mCherry^ is detected in the ventral nerve cord (vnc) and diffusely along peripheral nerves (arrows). **(D)** Adult *para*^*mCherry*^ nervous system stained for Para localization. Para^mCherry^ localizes prominently along segments of peripheral nerves (arrow) as they enter thoracic neuromeres. **(E)** Third instar larval nervous system stained with the pre-immune control. **(F,G)** Third instar larval nervous system (F) or adult brain (G) stained with affinity purified anti-Para antibodies. **(H)** Third instar larvae with the genotype [*para*^*mCherry*^; *OK371-Gal4, UAS-mCherry*^*dsRNA*^]. *para*^*mCherry*^ expression is suppressed in all glutamatergic neurons and thus Para^mCherry^ localization along axons of cholinergic sensory neurons becomes visible. **(I)** Third instar larvae with the genotype [*para*^*mCherry*^; *Chat-Gal4, UAS-mCherry*^*dsRNA*^]. Here expression of *para*^*mCherry*^ is suppressed in all cholinergic neurons which reveals Para localization in motor neurons. Note the prominent Para localization at the CNS/PNS transition point (arrowheads). **(J,K)** Ventral nerve cord of an adult fly with the genotype [*para*^*FlpTag-GFP*^; *Ok371-Gal4, UAS-flp*]. The boxed area is shown enlarged in (K). The arrowheads point to high density of Para. **(L)** Third instar larval *shaker*^*GFP*^ nervous system stained for GFP localization. Shaker is found in the neuropil (dashed area). **(M)** Third instar larval *shab*^*GFP*^ nervous system stained for GFP localization. Shab is distributed evenly along all peripheral axons. **(N)** Third instar larval *shal*^*GFP*^ nervous system stained for GFP localization. Shal localizes similar as Para on motor axons. Scale bars are 100 μm.

To independently assay Para localization, we generated antibodies against an epitope shared by all predicted Para isoforms. In western blots, anti-Para antibodies detect a band of the expected size (>250 kDa), which is shifted towards a higher molecular weight in protein extracts of homozygous *para*^*mCherry*^ animals (Figure S1D). Immunohistochemistry detects Para localization in control larvae but not in age matched *para* null mutant animals, further validating the specificity of the antibodies (Figure S1E). Staining of third instar larval filets revealed Para localization (Figure 1F) similar to what was noted for Para^mCherry^ localization (Figure 1C). Similarly, in the adult CNS, Para localization as detected by anti-Para antibodies matched the Para^mCherry^ signal (Figure 1D,G) Thus, we anticipate that endogenously mCherry-tagged Para protein reflects the wild typic Para localization.

To test a possible differential distribution of Para in either sensory or motor axons, we utilized RNAi to remove *mCherry* expression in *para*^*mCherry*^ heterozygous females. This leaves the wild type *para* allele intact and circumvents the early lethal phenotype associated with loss of *para*. Knockdown of *mCherry* expression in glutamatergicmotor neurons (Mahr and Aberle, 2006) reveals *para*^*mCherry*^ expression in cholinergic sensory neurons. Here, Para appears evenly localized along the entire length of the nerve (Figure 1H). In contrast, silencing *para*^*mCherry*^ in cholinergic neurons (Salvaterra and Kitamoto, 2001) reveals predominant localization of Para in an axonal segment of motor axons at the PNS/CNS boundary (Figure 1I). To differentiate between Para expression in motor and sensory neurons we also employed the FlpTag technique (Fendl et al., 2020). Upon Flp induced inversion of a GFP-encoding exon in adult motor neurons, strong Para localization is found in the part of the axon as it leaves the neuropil (Figure 1J,K).

### Differential localization of voltage-gated potassium channels

Several genes encode voltage-gated potassium channels. Using endogenously tagged proteins we find the Shaker potassium channel mostly in the synaptic neuropil regions (Figure 1L). The distribution of Shab resembles Para^mCherry^ localization in sensory axons (Figure 1H,M), whereas Shal localizes in a pattern similar to Para localization on motor axons (Figure 1I,N). Thus, Drosophila motor axons have an axonal segment resembling the vertebrate axon initial segment (AIS) harboring both voltage-gated sodium and potassium channels.

### High-resolution imaging reveals clustered localization of Para

To obtain a higher spatial resolution of Para distribution, we used structured illumination microscopy. Para localization distal to the AIS is found in a clustered arrangement with a spacing of 0.6-0.8 μm (Figure 2A-B’). Flp expression only in one motor neuron in each larval hemineuromer using *94G06-Gal4* (Pérez-Moreno and O’Kane, 2019), allows labeling of single axons when using the FlpTag technique. Weak expression is noted around the nucleus (Figure 2C, arrow). Flanking the strong expression along the AIS, a clustered localization of Para^GFP^ was apparent which, however, was too faint to allow high resolution microscopy (Figure 2D, arrowheads). To exclude that cluster formation is due to the fluorescence protein moiety, we performed anti-Para immunohistochemistry on primary Drosophila neural cells in culture, where axons form small fascicles with few accompanying glial cells. When such cultures are stained for Para distribution, we find Para channels localized in small clusters with a spacing of about 0.6-0.8 μm (Figure 2E,F). In conclusion, next to an axon initial segment that likely serves as a spike initiation zone (Günay et al., 2015), Para channels are organized in a clustered arrangement, supporting the notion that micro-saltatory conductance contributes to an increase in axonal conductance velocity.

**Figure 2.**
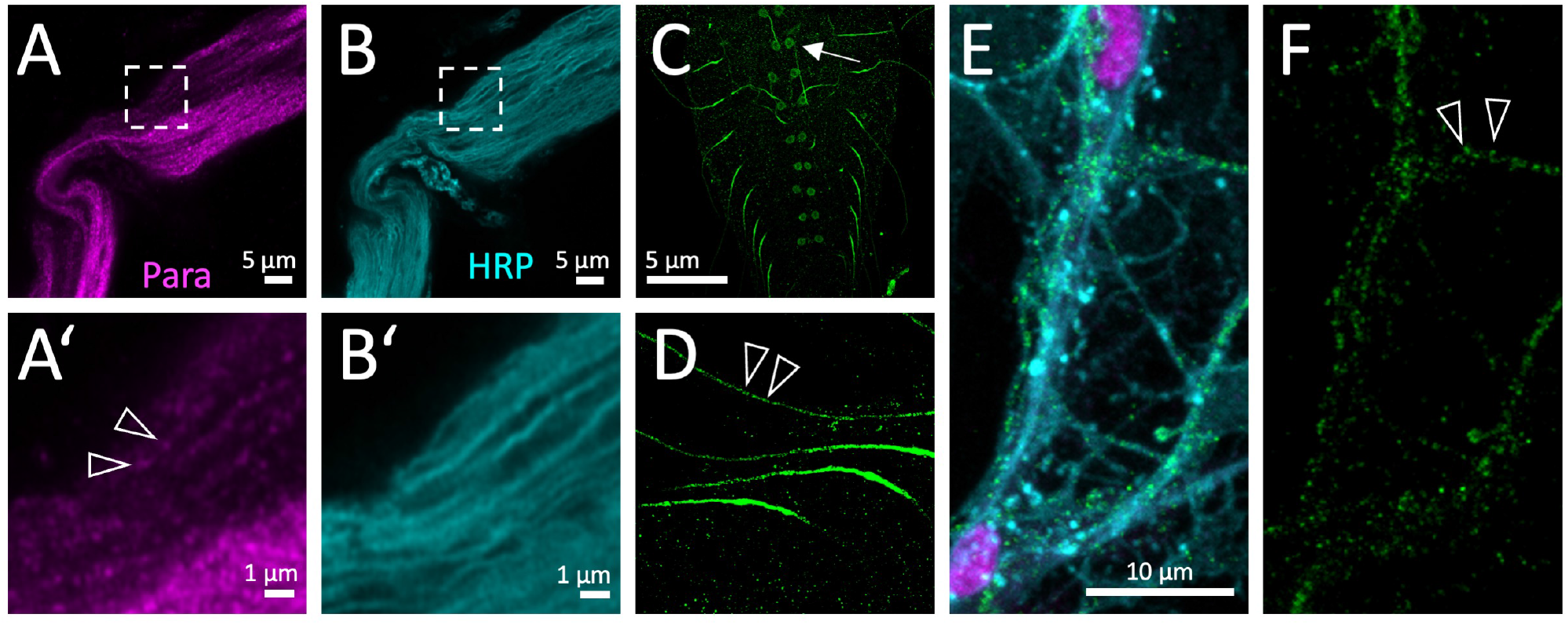
Clustered localization of Para. **(A)** High resolution Airyscan analysis of Para^mCherry^ and **(B)** HRP localization in an adult nerve. The boxed area is shown in higher magnification below (*A’,B’*). Note the clustered appearance of Para^mCherry^, clusters are about 0.6-0.8 μm apart (arrowheads). **(C)** Ventral nerve cord of a third instar larva with the genotype [*para*^*FlpTag-GFP*^; *94G06-Gal4, UAS-flp*]. The arrow points to a neuronal cell body. **(D)** Higher magnification of single Para^GFP^ expressing axons. Note the dotted arrangement of Para^GFP^ along the motor axon (arrowheads). **(E,F)** Primary wild type neurons cultured for 7 days stained for Repo (magenta) to label glial nuclei, HRP (cyan) to label neuronal cell membranes and anti-Para antibodies (green) (E). The Para protein localizes in a dotted fashion (FScale bars are as indicated.

### A lacunar system localizes around axonal segments rich in Para

In vertebrates, most voltage-gated ion channels are found at the axon initial segment (AIS) (Eshed-Eisenbach and Peles, 2019). To determine the distribution of Para on the subcellular level, we integrated an Apex2 encoding exon in the *para* locus which allows generating local osmiophilic diaminobenzidine (DAB) precipitates that are detectable in the electron microscope (Lam et al., 2015). This resulted in a weak hypomorphic *para* allele (Figure 1A,B). In adult flies, Para^Apex2^ is expressed in sufficient intensity to be detected along peripheral axons using the electron microscope (Figure 3A, white arrowheads). In segments next to the CNS/PNS boundary revealed differential DAB precipitates along single large axons (Figure 3B-D). In longitudinal sections of Para-rich axon segments we determined the staining intensity along the plasma membrane by using Fiji (Figure 3E,F). Here, the modulation of staining intensity is apparent with a spacing resembling what we determined by confocal microscopy (2A’,F, Figure 3E,F).

**Figure 3.**
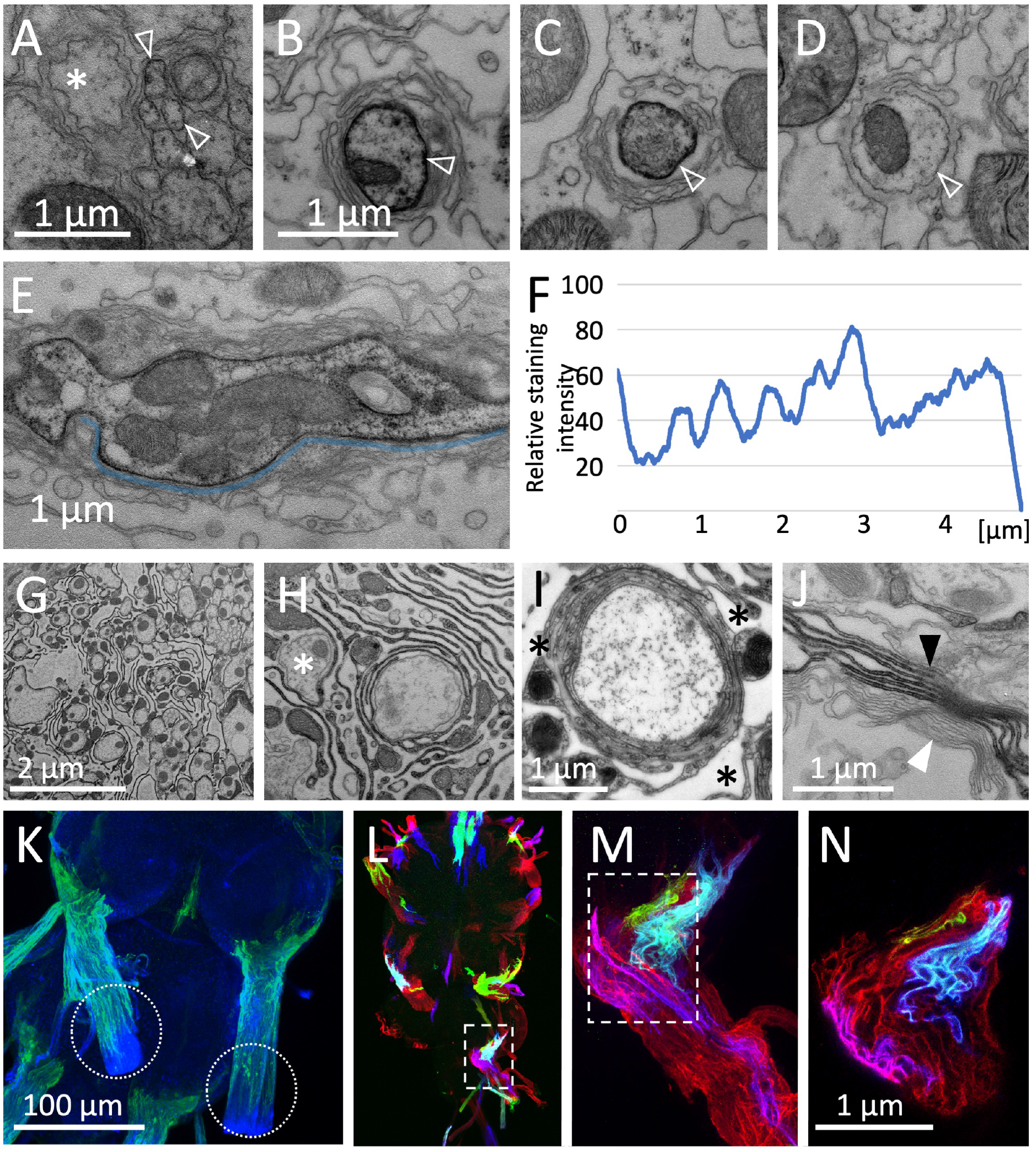
Glial lacunar system surrounds the AIS. (**A)** Weak Para expression can be detected on *para*^*Apex2*^ expressing small axons (arrowheads) running in fascicles within the nerve. **(B-D)** Semi-serial cross-sections through the nerve. The same axon is shown in all images. Distance between individual sections (B,C) 15μm, distance between (C,D) 6.5 μm. Note the intense labeling of the axonal membrane is changing between the different sections. **(E)** Longitudinal section of a *para*^*Apex2*^ expressing axon. To determine the staining intensity along the membrane (above the blue line) a corresponding ROI was defined and quantified using Fiji. **(F)** Quantification of the staining intensity of the membrane stretch shown in (E). Note the regular increase in staining intensity every 0.6-0.8 μm. **(G,H,J)** Apex2 expression directed by *75H03-Gal4*. **(G)** Overview of the lacunar organization at the CNS/PNS boundary. (**H)** Lacunar structures are largely formed by the tract glia. The asterisk highlights an axon with only one direct glial wrap. **(I)** High pressure freezing preparation showing a single axon covered by myelin-like membrane sheets in a lacunar area (asterisks). **(J)** Darkly stained tract glia membrane stacks (black arrowhead) can be found next to unlabeled membrane stacks (white arrowhead) suggesting that myelin-like structures can be derived from both, central and peripheral wrapping glial cells. **(K)** *75H03-Gal4* directed expression of GFP labels the ensheathing/wrapping or tract glia. Note that GFP expression ends proximal to the dissection cut (white dashed circles). **(L-N)** MCFO2 analysis of the *nrv2-Gal4, R90C03-Gal80* positive wrapping glia. Note that glial cells tile the nerve roots with no gaps in between. Scale bars are as indicated.

Interestingly, highest levels of Para localization coincide with a mesh-like glial organization around large caliber axons, that resembles a lacunar system (Figure 3G,H) (Wigglesworth, 1960). The glial lacunar system is characterized by intensive formation of glial processes around a few axons (Figure 3H, asterisk). In addition we find axons with multiple wrapping (Figure 3I). Next we determined which glial cell type form these lacunar structure. In the larva, peripheral wrapping glia express the *nrv2-Gal4 90C03-Gal80* driver (Kottmeier *et al*., 2020; Matzat *et al*., 2015; Stork *et al*., 2008) whereas central ensheathing/wrapping glia express the *83E12-Gal4* driver (Peco et al., 2016; Pogodalla et al., 2021) (Figure S1A,B). In adults, a specialized group of glia is found at the CNS/PNS boundary, called tract glia (Kremer et al., 2017) (Figure S2E), similar to what is known from the vertebrate nervous system (Fontenas and Kucenas, 2017; Kucenas et al., 2008; Kucenas et al., 2009). The position of the lacunae coincides with the location of the tract glial cells (Kremer *et al*., 2017)(Figure 3K). They express *75H03-Gal4, 83E12-Gal4* and the *nrv2-Gal4 90C03-Gal80* driver (Figure S2). Multi color flipout (MCFO2) labeling experiments (Nern et al., 2015) indicate tiling of these glial cells along the nerve with no overlap and no spaces in between individual glial cells (Figure 3L-N).

Apex2 expression experiments confirmed, that most of the lacunar system is indeed generated by wrapping and tract glial cell processes (Figure 3,G-J, black and white arrowheads, Figures S2,3). Thus, in large caliber motor axons, most of the Para voltage-gated sodium channel appears to be positioned close to the lacunar system, which had been speculated to serve as an extracellular ion reservoir needed for sustained generation of action potentials (Chandra and Singh, 1983; Leech and Swales, 1987; Maddrell and Treherne, 1967; Treherne and Schofield, 1981; Van Harreveld et al., 1969; Wigglesworth, 1960).

### Myelin in the leg nerve is found close to the CNS

In vertebrates, clustering of voltage-gated ion channels occurs on the edges of myelinated axonal segments (internodes) (Arancibia-Cárcamo *et al*., 2017; Castelfranco and Hartline, 2015; Cohen *et al*., 2019; Dutta *et al*., 2018; Eshed-Eisenbach and Peles, 2019). Myelin not only participates in positioning of voltage-gated ion channels but also increases electric insulation and thus contributes to a faster conductance velocity. In Drosophila highest conductance velocity is likely to be required during fast and well-tuned locomotion in adults. Thus, we focused our analysis on adult leg nerves. Most of the 760 axons (Figure S1F) within an adult leg run in a single large nerve that exit the CNS at well-defined positions (Figure 4A,B). Here, axons are organized in zones depending on their diameter (Figure 4C,D). At the position of the femur, large axons are covered by a single glial sheet (Figure 4C). At the coxa, close to the CNS, we noted that large diameter axons were engulfed by several glial membrane sheets (Figure 4D, asterisk). Up to 15 flat glial membrane sheets with a thickness of about 28 nm are found along larger axons (Figure 4E,F, Figure S1G), whereas small diameter axons are engulfed in a bundle by a single glial wrap (Figure 3G). Axons with an intermediate diameter show individual glial wrapping with a single or very few glial sheets (Figure 4E-G).

**Figure 4.**
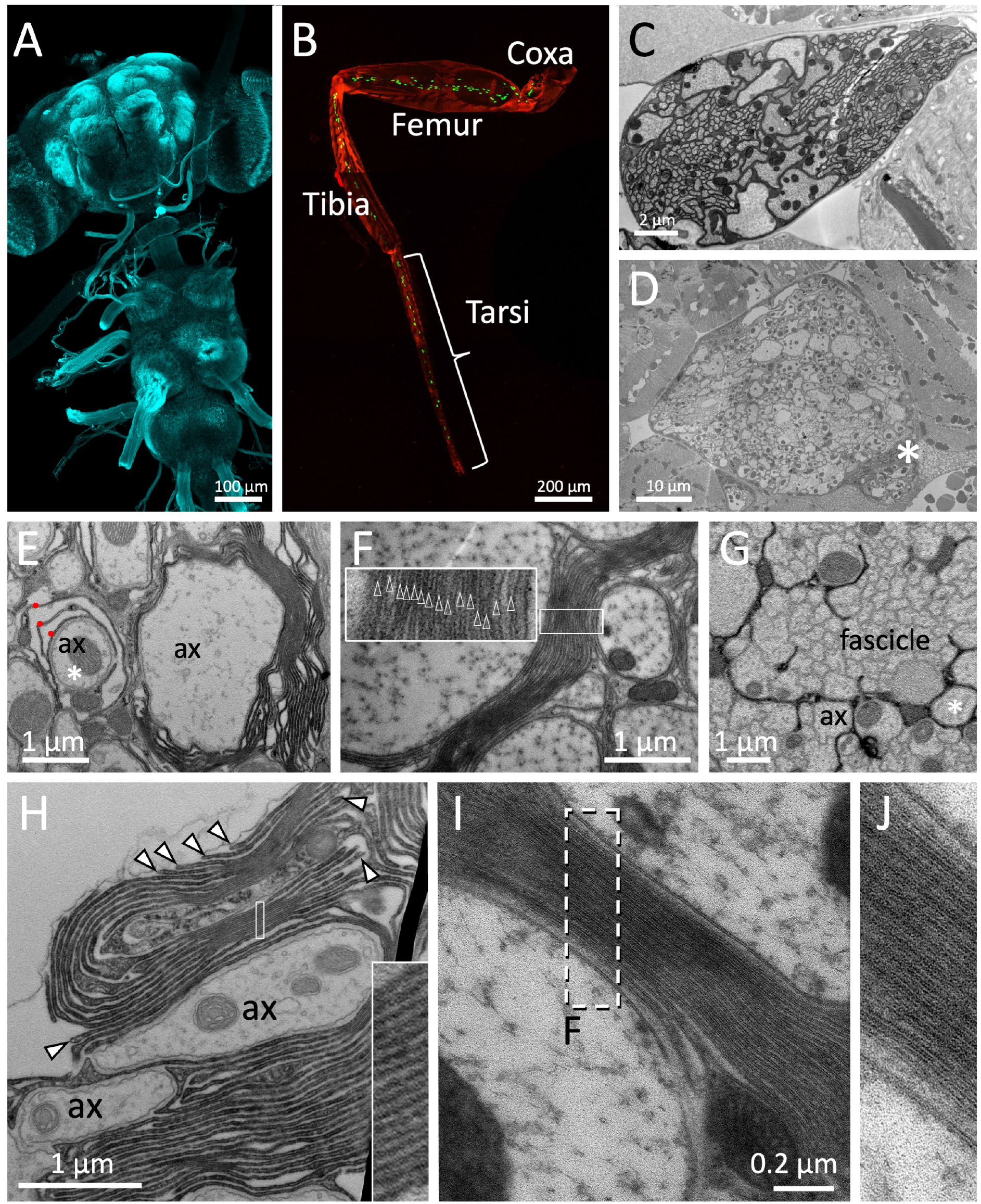
Drosophila wrapping glia form myelin. **(A)** Overview of an adult central nervous system, stained for HRP (cyan). **(B)** Drosophila leg with wrapping glial nuclei in green, genotype [*nrv2-Gal4, UAS-lamGFP*]. **(C)** Electron microscopic section at the level of the femur. **(D)** Electron microscopic section at the level of the coxa. Note the increased amount of glial membranes around the large caliber axons which is indicated by the thick grey material around the axons (asterisk). **(E,G,H)** Glial cell processes are stained by the presence of Apex2 which generates an osmiophilic DAB precipitate [*75H03-Gal4, UAS-Myr-Flag-Apex2-NES*]. **(E)** Large caliber axons are surrounded by glial membrane stacks. The asterisk denotes an axon engulfed by a few glial wraps (red dots). ax: axon. **(F)** Up to 15 densely packed membrane sheets are found (see inlay for enlargement). **(G)** In the leg nerve, small axons are engulfed as fascicles and are not individually wrapped. The asterisk denotes an axon engulfed by a single glial wrap. ax: axon. **(H)** Note the bulged appearance of the growing tip of the glial cell processes that form the myelin-like structures (arrowheads). The inlay shows a highly organized membrane stacking. **(I,J)** High pressure freezing preparation of prefixed samples to reduce tissue preparation artifacts. Note the compact formation of membrane layers.

### Myelin is formed by peripheral wrapping glia

To determine whether myelin-like structures are made by glia, we generated flies harboring a *UAS-Myr-Flag-Apex2-NES* transgene, Following expression of Apex2^Myr^ in *75H03-Gal4* expressing glial cells, processes can be clearly visualized (Figure 4G,H, Figure S3). In some areas close to the CNS we noted almost compacted glial membrane sheets (Figure 4H, inlay boxed area). We performed additional high pressure freezing of pre-fixed samples to optimize tissue preservation (Sosinsky et al., 2008). In such specimens compact stackings of thin glial membrane sheets can be detected, too (Figure 4I,J). Thus, Drosophila glia is able to form myelin at the CNS/PNS junction.

One notable difference to vertebrate myelin is the lack of spiral growth of wrapping glial processes, although it can be detected occasionally (Figure S3A, Figure S4). Drosophila glial membrane stacks are formed by extensive membrane folding close to the axon providing the disadvantage that axons are not entirely insulated (Figure 4H, Figure S3). An additional unique feature of vertebrate myelin is its compact organization which is mediated by the myelin basic protein (MBP) (Nave and Werner, 2021). In contrast to vertebrate myelin where extra- and intercellular space is removed, fly myelin-like structures only show an irregular compaction of the extracellular space.

### Para localization depends on wrapping glial cells

To next test whether wrapping glial cells control positioning of voltage-gated ion channels, we ablated either peripheral glia or central wrapping glia by the proapoptotic gene *hid* (Kottmeier *et al*., 2020; Pogodalla *et al*., 2021) and assayed the distribution of Para. Removal of the CNS specific ensheathing glia did not affect Para localization in the larval nervous system (Figure S5A-E). In contrast, upon ablation of the peripheral wrapping glia a marked change in Para protein localization becomes obvious (Figure 5A-B’). In control larvae, anti-Para antibodies detect only a weak labeling of segmental nerves. In contrast, all nerves are intensely decorated with Para in wrapping glia ablated larvae. Further qRT-PCR experiments demonstrate a two-fold increase of *para* mRNA levels (Figure 5C).

**Figure 5.**
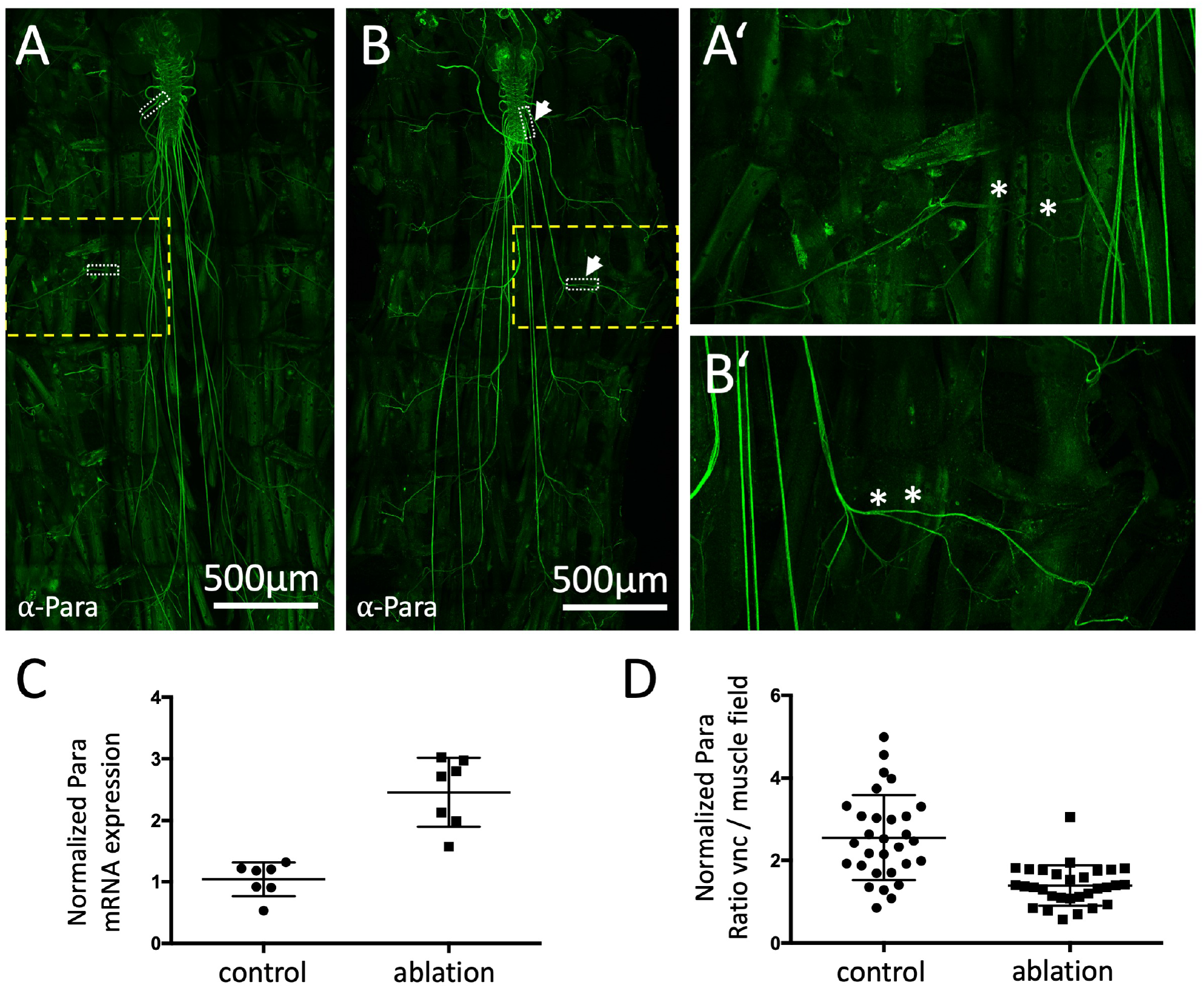
Localization of the voltage gated sodium channel depends on glia. **((A,A’)** Third instar larval filet preparation with the genotype [*nrv2-Gal4, UAS-CD8-GFP; R90C03-Gal80*] showing the localization of Para as detected using the anti-Para antibody in a control larva. **(B,B’)** Third instar larval filet preparation with the genotype [*nrv2-Gal4, UAS-hid; R90C03-Gal80*] showing the localization of Para as detected using the anti-Para antibody in a wrapping glia ablated larva. The white dashed boxes were used for quantification of Para fluorescence intensity in the CNS/PNS transition zone in relation to its expression in the muscle field area. The yellow boxed areas are shown in higher magnification (A’,B’). Note the increased localization of Para along the peripheral nerve at the level of the muscle field (asterisks). Scale bars are as indicated. **(C)** Quantification of *para* mRNA expression using qRT-PCR in control and wrapping glia ablated larvae (n = 7, with 15-20 brains each). Note, the increase in para mRNA expression upon wrapping glia ablation. **(D)** Quantification of relative Para localization in the CNS/PNS transition area and the muscle field area in control and wrapping glia ablated larvae (n = 10 larval filets, with 30 nerves, p = 0,0047, unpaired t-test). Note, Para distributes evenly along the axon in the absence of wrapping glia.

Finally, we not only noted an upregulation of *para* mRNA and protein expression, but also a redistribution along axons upon glial cell ablation. Whereas in wild type control larvae, 2.5 times more Para protein is found at the CNS/PNS transition zone compared to nerve segments on the muscle field, an almost even distribution is noted in glia ablated larvae (Figure 5D). Interestingly, ablation of central or peripheral wrapping glial cells does not affect the distribution of the voltage gated potassium channel Shal (Figure S6).

Taken together, even in the small insect *Drosophila melanogaster*, myelin-like structures are formed (Figure 6). They flank specific lacunar regions formed by glial cell processes that possibly provide a large extracellular ion reservoir needed for the generation of action potentials (Figure 6). Para voltage-gated sodium channels are differentially localized along sensory and motor neurons. In motor neurons Para localization is enriched in axonal segments within the lacunar system, and its normal expression and its localization is dependent on the presence of wrapping glial cell processes. This suggests a signaling pathway from glia to the regulation of *para* transcription.

**Figure 6.**
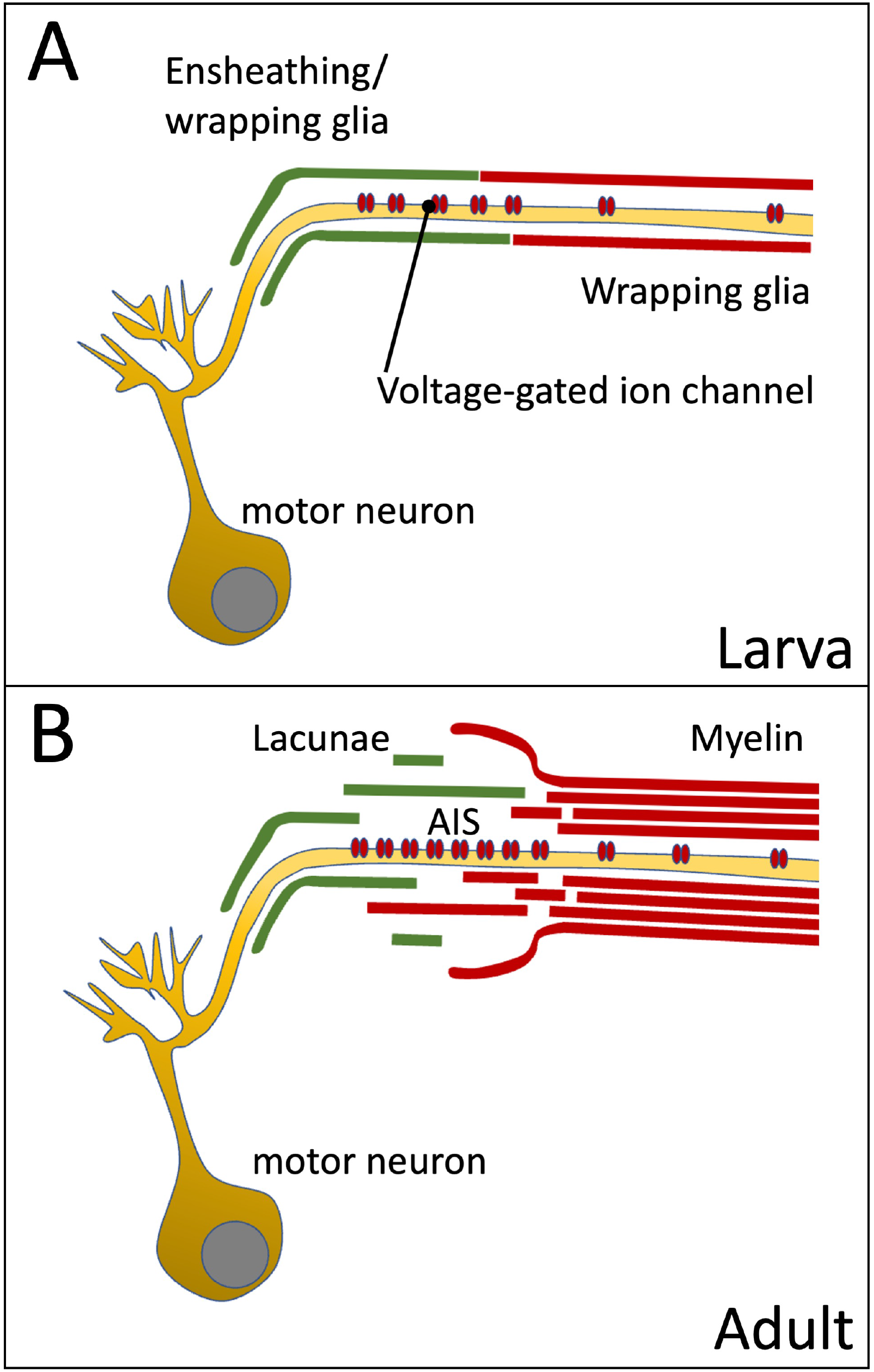
Organization of the axon initial segment in Drosophila motor axons Voltage gated sodium channels are preferentially positioned at the axon initial segment (AIS) of the motor axon. **(A)** In the larval nervous system positioning is mediated by the peripheral wrapping glia. **(B)** In adults these cells form myelin-like structures, which fray out in the lacunae which represent a reservoir possibly needed for ion homeostasis during sustained action potential generation.

## Discussion

In the vertebrate nervous system, saltatory conductance allows very fast spreading of information. This requires localized distribution of voltage-gated ion channels and concomitantly, the formation of the myelin sheath. The evolution of this complex structure is unclear. Here, we report glial-dependent localization of voltage-gated ion channels at an AIS-like domain of peripheral Drosophila larval motor axons. As more channels accumulate in adults, a lacunar system and adjacent myelin-like structures are formed by wrapping glia.

In myelinated axons of vertebrates, voltage-gated Na^+^ and K^+^ channels are clustered at the AIS and the nodes of Ranvier (Amor et al., 2014; Freeman *et al*., 2016; Nelson and Jenkins, 2017). The AIS is the spike initiation zone located distal to the soma and distal to the dendrite branching point of invertebrate neurons (Günay *et al*., 2015). AIS-like segments have been previously postulated for Drosophila neurons due to the localization of a giant ankyrin, since in vertebrates the AIS formation depends on this scaffolding protein (Dubessy et al., 2019; Freeman et al., 2015; Jegla et al., 2016; Trunova et al., 2011). Moreover, recent modeling approaches at the example of the pioneering aCC motor neuron predicted the localization of voltage-gated ion channels at the CNS/PNS boundary (Günay *et al*., 2015), which very well matches the localization of the voltage-gated ion channels Para and Shal, as described here. Interestingly, in Drosophila *para* mRNA expression as well as Para protein localization depend on the presence of peripheral wrapping glia. In glia ablated nerves, Para localizes evenly along the entire axonal membrane. This loss of a clustered distribution may contribute to the pronounced reduction in axonal conductance velocity noted earlier in such glia ablated animals (Kottmeier *et al*., 2020). In addition, we found an increased *para* mRNA expression. How glial cells control Para localization and how this is then transduced to an increased expression of *para* remains to be further studied. Possibly, positioning of voltage-gated ion channels may require glial secreted proteins (Yuan and Ganetzky, 1999).

In the adult nervous system, the AIS-like domain is embedded in glial lacunar regions formed by wrapping glial cell processes. The increased expression of Para within the AIS-like segments of adult brains (Ravenscroft *et al*., 2020) is expected to generate strong ephaptic coupling forces (Kottmeier *et al*., 2020; Rey et al., 2020). These are caused by ion flux through open channels which generate an electric field that is able to influence the gating of ion channels in closely neighboring axons (Arvanitaki, 1942; Krnjevic, 1986; Rasminsky, 1980). Ephaptic coupling helps to synchronize firing axons (Anastassiou and Koch, 2015; Anastassiou et al., 2011; Han et al., 2018; Shneider and Pekker, 2015), but is also detrimental to the precision of neuronal signaling in closely apposed axons (Arvanitaki, 1942; Kottmeier *et al*., 2020). This is counteracted by the glial lacunar system, that spatially separates axons and adds more levels of wrapping. Furthermore, it was postulated that the lacunar system provides a large extracellular ion reservoir. This would allow sustained neuronal activity required for action potential generation, which would not be easily possible given the small interstitial fluid volume normally found between two cells. The lacunar structures than appear to collapse to form compact myelin-like membrane sheets. From here on distally, ion channel density decreases. Concomitantly, the need for a large ion reservoir decreases favoring the formation of myelin-like structures.

A hallmark of vertebrate myelin is the spiral growth of the insulating glial membrane. This is generally not observed in large fly nerves where glial membrane sheets rather fold back than spirally grow around a single axon. Compared to myelinated vertebrate axons, this provides the disadvantage that axons are not entirely insulated. However, spiral growth can be seen in small nerves where less extensive wrapping is noted. An additional unique feature of vertebrate myelin is its compact organization which is mediated by the myelin basic protein (MBP) (Nave and Werner, 2021). In contrast to vertebrate myelin where extra- and intercellular space is removed, fly myelin-like structures only show a compaction of the extracellular space, which is expected to increase resistance as the number of freely moving ions is diminished. A fully compact myelin state would require MBP-like proteins which have not been identified in the fly genome.

In conclusion, the evolution of myelin appears reflected in the different developmental stages of Drosophila. First, voltage gated ion channels are clustered at the AIS with the help of Drosophila glia (Figure S6A). Second, upon increased expression of such ion channels in the adult nervous system, an ion reservoir is formed by the lacunar system. The collapse of glial processes in the non-lacunar regions then provides the basis of myelin formation. In the future, it will be interesting to identify glial derived signals that ensure channel positioning and determine how neuronal signaling adjusts channel expression and triggers formation of myelin.

## Materials

### Drosophila genetics

All fly stocks were raised and kept at room temperature on standard Drosophila food. All crosses were raised at 25 °C. The following fly lines were used:

| Fly stock | Source |
| --- | --- |
| **MiMIC fly lines** |  |
| *y(R Caré et al.) w[\*] Mi{MIC}MI08578a*<br>*Mi{MIC}MI08578b* | BDSC51078 |
| *y(R Caré et al.) w[\*]*<br>*Mi{FlpStop}para[MI08578-FlpStop.D]/FM7c* | BDSC 67680 |
| *Para-mCherry* | This study |
| *Para-Apex2* | This study |
| *Para-FlpTag GFP/Fm7i* | (Fendl *et al.*, 2020) |
| *y(R Caré et al.) w[\*]; Mi{PT-GFSTF.1}Shal[MI00446-GFSTF.1]* | BDSC 60149 |
| *y(R Caré et al.) w[\*] Mi{PT-GFSTF.2}Sh[MI10885-GFSTF.2]/FM7j, B(R Caré et al.)* | BDSC 59423 |
| *y(R Caré et al.) w[\*]; Mi{PT-GFSTF.1}Shab[MI00848-GFSTF.1]/TM6C, Sb(R Caré et al.) Tb(R Caré et al.)* | BDSC 60514 |

| UAS lines |  |
| --- | --- |
| *y(R Caré et al.) sc[\*] v(R Caré et al.) sev[21];*<br>*P{y[+t7.7] v[+t1.8]=VALIUM20-mCherry}attP2* | BDSC 35785 |
| *UAS-CD8GFP II; R90C03Gal80 III* | (Kottmeier et al., 2020) |
| *UAS-Hid/CyOw; R90C03Gal80 III* | (Kottmeier et al., 2020) |
| *UAS-lacZ NLS* | Kindly provided by Y. Hiromi |
| *UAS-lambda-Htl* | (Gisselbrecht et al., 1996) |
| *UAS-htl<sup>DN</sup> II* | BDSC 5366 |
| *UAS-CD8GFP II* | BDSC 5137 |
| *UAS-CD8mCherry II* | BDSC 27391 |
| *UAS-Hid* | (Igaki et al., 2000) |
| *UAS-Flp* | BDSC 4539 |
| *UAS-Myr-Flag-APEX2-NES<sup>86Fb</sup> III* | This study |
| *pBPhsFLP2::pEST / I ; ; UAS HA, FLAG, V5,*<br>*OLLAS / III* | (Nern et al., 2015) |
| Gal4 lines |  |
| *ChAT-Gal4* | (Salvaterra and Kitamoto, 2001) |
| *OK371-Gal4* | (Mahr and Aberle, 2006) |
| *GMR94G06Gal4 III* | BDSC 40701 |
| *Nrv2-Gal4 II* |  |
| *Nrv2Gal4 II; R90C03Gal80 III* | (Kottmeier et al., 2020) |
| *Nrv2Gal4 II; R90C03Gal80, UAS-CD8Cherry III* | (Kottmeier *et al.*, 2020) |
| *GMR83E12_AD II; Repo4.3_DBD III* | (Bittern *et al.*, 2021) |
| *GMR75H03-Gal4 III* | BDSC 39908 |
| **lexAop lines** |  |
| *lexAop-Flp (III)* | BDSC 55819 |
| **lexA lines** |  |
| *w[\*]; Tl{2A-lexA::GAD}VGlut[2A-lexA]/CyO* | BDSC 84442 |
| *para<sup>ST76</sup>* | BDSC 26701 |
| Oregon R | BDSC 5 |

## Methods

To determine temperature sensitivity, five 3-days old male and female flies were transferred to an empty vial with a foam plug. The vials were incubated in a water bath at 42°C for 1 min and then placed at room temperature. Flies were monitored every 15 secs for 5 min. At least 100 males and females for each genotype were tested.

For MCFO experiments early, white pupae were collected, put in a fresh vial and heat shocked at 37 °C for 1 hour. Pupae were placed back to 25 °C and dissected a few days after hatching. To generate *para*^*mCherry*^ flies, we injected pBS-KS-attB1-2-PT-SA-SD-0-mCherry as described (Venken *et al*., 2011). To generate *para*^*Apex2*^ flies, we inserted the *apex2* coding sequence (addgene #49386, using the primers AAGGATCCGGAAAGTCTTACCCAACTGT and AAGGATCCGGCATCAGCAAACCCAAG) into pBS-KS-attB1-2-PT-SA-SD-0-mCherry (DGRC#1299) after mCherry was deleted using appropriate restriction enzymes. pBS-KS-attB1-2-PT-SA-SD-0-Apex2 was used to establish a *para*^*Apex2*^ as described (Venken *et al*., 2011). Flies were tested via single-fly PCR. UAS-Myr-Flag-Apex2 flies were generated by cloning Apex2 using the primers CACCgactacaaggatgacgacgataa and cagggtcaggcgctcc into pUAST_Myr_rfA_attB

### Western blot analysis

10 adult fly heads were homogenized in 50 μl RIPA buffer on ice. They were centrifuged at 4°C for 20 min at 13.000 rpm. The supernatant was mixed with 5x reducing Lämmli buffer and incubated for 5 min at 65°C. 15 μl of the samples were separated to an 8% SDS-gel and subsequently blotted onto a PVDF membrane (Amersham Hybond-P PVDF Membrane, GE Healthcare). Anti-Para antibodies were generated against the following N-terminal sequence (CAEHEKQKELERKRAEGE), affinity purified, and were used in a 1/1,000 dilution. Experiments were repeated three times.

### Cell culture

Primary neural cell culture was preformed as described (Prokop et al., 2012). In brief, 3-5 stage11 embryos were collected, chemically dechorionized and homogenized in 100 μl dispersion medium. Following sedimentation for 5 min at 600 g, cells were resuspended in 30 μl culture medium and applied to a glass bottom chamber (MatTek), sealed with a ConA coated coverslip. Cultures were grown for 5-7 days. Experiments were repeated three times.

### qPCR

RNA was isolated from dissected larval brains using the RNeasy mini kit (Qiagen) and cDNA was synthesised using Quantitect Reverse Transcription Kit (Qiagen) according to manufacturer’s instructions. qPCR for all samples was performed using a Taqman gene expression assay (Life technologies) in a StepOne Real-Time PCR System (Thermofisher, para: Dm01813740_m1, RPL32: Dm02151827_g1). RPL32 was used as a housekeeping gene. Expression levels of Para were normalized to RPL32.

### Immunohistochemistry

Larval Filets: L3 wandering larvae were collected in PBS on ice. Larvae were placed on a silicon pad and attached with two needles at both ends, with the dorsal side facing up. They were cut with a fine scissor at the posterior end. Following opening with a long cut from the posterior to the anterior end the tissue was stretched and attached to the silicon pad with additional 4-6 needles. Gut, fat body and trachea were removed. Adult brains: Adult flies were anesthetised with CO_2_ and were dipped into 70% ethanol. The head capsule was cut open with fine scissors and the tissue surrounding the brain removed with forceps. Legs and wings were cut off and the thorax opened at the dorsal side. The ventral nerve cord was carefully freed from the tissue. For fixation, dissected samples were either covered for 3 min with Bouin’s solution or for 20 min with 4% PFA in PBS. Following washing with PBT samples were incubated for 1 h in 10% goat serum in PBT. Primary antibody incubation was at 4°C followed. The following antibodies were used: anti-Para N-term, this study; anti-dsRed (Takara), anti-GFP (Abcam, Invitrogen), anti-Rumpel (Yildirim et al., 2022), anti-Repo (Hybridoma bank), rabbit α-V5 (1:500, Sigma Aldrich), mouse α-HA (1:1000, Covance), rat α-Flag (1:200, Novus biologicals). The appropriate secondary antibodies (Thermofisher) were incubated for 3 hrs at RT. The tissues were covered with Vectashield mounting solution (Vector Laboratories) and stored at 4°C until imaging using a LSM880 (Carl Zeiss AG). All stainings were repeated >5 times.

### High Pressure Freezing

3 weeks old female flies were used with head, legs and tip of abdomen removed. Following fixation in 4% FA in 0.1M PHEM in a mild vacuum (−200 mbar), at RT for 45 min and 3 washes in 0.1 PHEM, the tissue was embedded in 3% low melting agarose for vibratome sectioning (Leica, VTS1200S). Samples were cut in PBS into 200 μm thick cross sections with 1mm/sec, 1.25mm amplitude. and were placed into lecithin coated 6 mm planchettes, filled with 20% PVP in 0.1M PHEM and high pressure frozen (Leica, HPM100). 7 specimens were sectioned. Freeze substitution was performed in 1 %OsO_4_, 0.2%glutaraldehyde, 3% water in acetone at -90°C and stepwise dehydrated over 3days. Samples were embedded in mixtures of acetone and epon.

### DAB Staining and electron microscopy

Flies were injected with 4% formaldehyde (FA) in 0.1 M HEPES buffer and fixed at room temperature for 45 min. Following washes and incubation in 20 mM glycine in 0.1 M HEPES, samples were incubated in 0.05 % DAB in 0.1 M HEPES at room temperature for 40 min. 0.03% H_2_O_2_ was added and the reaction was stopped after 5-10 min. The tissue was then fixed in 4% FA and 0.2% glutardialdehyde in 0.1 M HEPES at RT for 3 h. After 3 times rinsing the tissue was fixed in 4% FA at room temperature overnight. The FA was replaced by 2% OsO_4_ in 0.1 M HEPES for 1 h on ice (dark). Uranyl acetate staining was performed *en bloque* using a 2% solution in H_2_O for 30 min (dark). Following an EtOH series (50%, 70%, 80%, 90% and 96%) on ice for 3 min each step, final dehydration was done at room temperature with 2x 100 % EtOH for 15 min and 2 times propylene oxide for 30 min. Grids of high pressure frozen samples were additionally counterstained with uranyl acetate and lead citrate. Following slow Epon infiltration specimens were embedded in flat molds and polymerized at 60 °C for 2 days.

6 specimens from 3 different fixation experiments were sectioned. Ultrathin sections were cut using a 35° ultra knife (Diatome) and collected in formvar coated one slot copper grids. For imaging a Zeiss TEM 900 at 80 kV in combination with a Morada camera (EMSIS, Münster, Germany) operated by the software iTEM. Image processing was done using Adobe Photoshop and Fiji. Ultrathin sections of high pressure frozen samples were examined at a Tecnai 12 biotwin (Thermo Fisher Scientific) and imaged with a 2K CCD veleta camera (EMSIS, Münster, Germany).

## Acknowledgements

We are grateful to all our colleagues for many discussions and P. Deing, K. Krukkert, K. Mildner and E. Naffin for excellent technical assistance. B. Zalc and K.A Nave for critical reading of the manuscript and many thoughtful suggestions. This work was supported by the Deutsche Forschungsgemeinschaft through funds to C.K. (SFB 1348, B5, Kl 588 / 29).

## Author contribution

H.O. analyzed the *para* gene and S.R. conducted the electron microscopic analyses. F.M. generated the Apex2 encoding exon. D.Z. designed the high pressure freezing experiments. C.K. wrote the first version of the manuscript and S.R., H.O., and C.K. discussed and revised the text.

## Data and materials availability

All Drosophila strains reported are available upon request to C.K..

## Supplementary Figures

**Figure S1.**
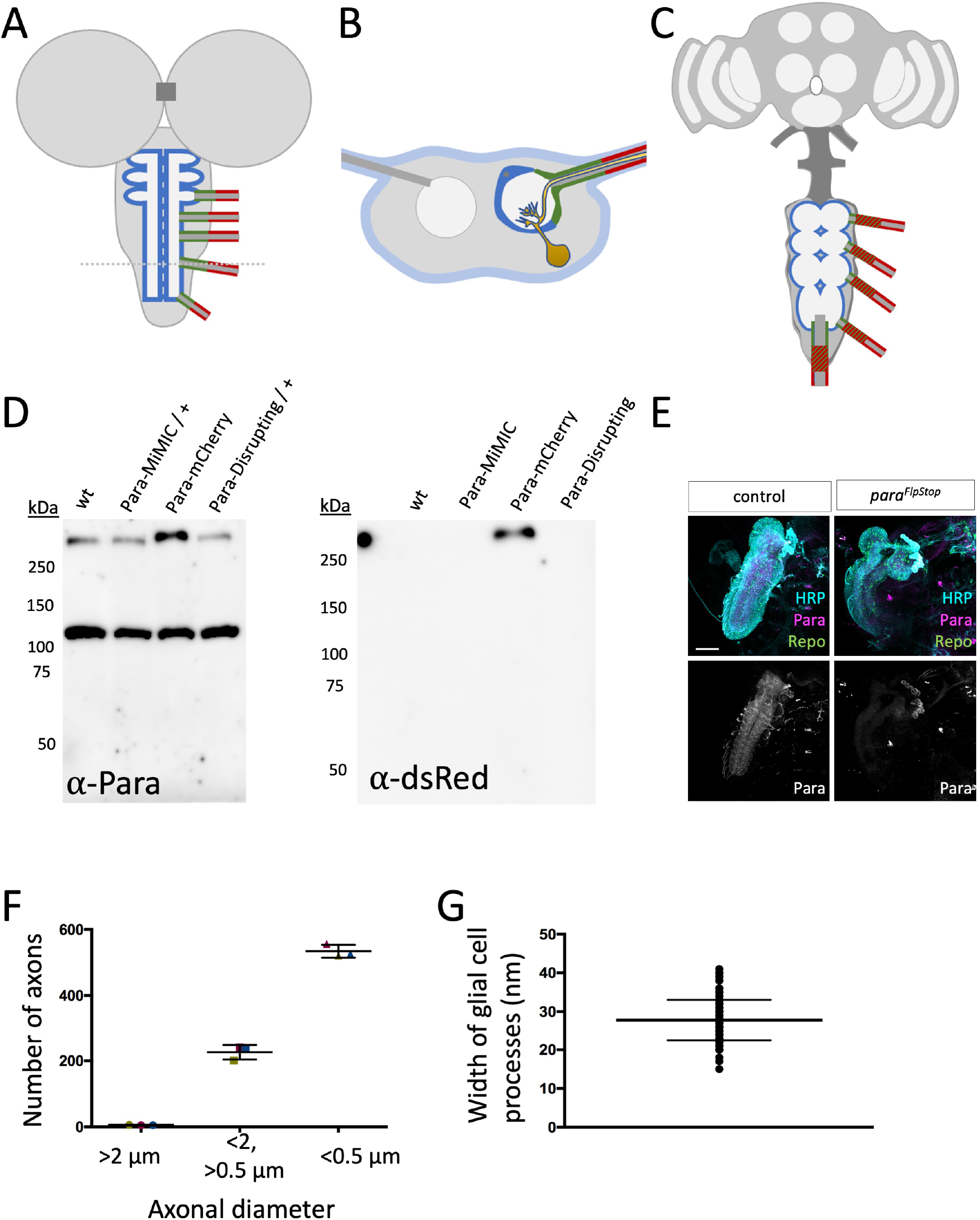
**(A-C)** Schematic representation of the larval (A,B) and the adult Drosophila nervous system (C). The ensheathing glia is labelled in blue, the ensheathing/wrapping glia is labelled in red, the wrapping glia is shown in green. The tract glia of the adult nervous system is shown in green and red stripes. The tract glia likely corresponds to the ensheathing/wrapping glia but the exact lineage relationship is not known. **(D)** Western blot of protein lysates of adult heads. Purified anti-Para antibodies detect a band of 105 kDa and a band of >250 kDa in size. The size of the >250 kDa protein band increases in *para*^*mCherry*^ heads compared to wild type control as well as *para*^*MiMIC*^ heads, indicating that this band corresponds to the Para protein. Note that elevated levels of the endogenous Para::mCherry fusion protein are detected. Anti-dsRed antibodies detect only the Para^mCherry^ fusion protein. **(E)** Para antisera detect a protein in dissected 24 hours old wild type animals. No protein is detected in dissected age-matched para mutant animals. **(F)** About 760 axons innervate the leg. The majority is smaller than 0.5 μm in diameter, very few ones are larger than 2 μm. **(G)** The width of glial cell processes is about 28 nm and very regular.

**Figure S2.**
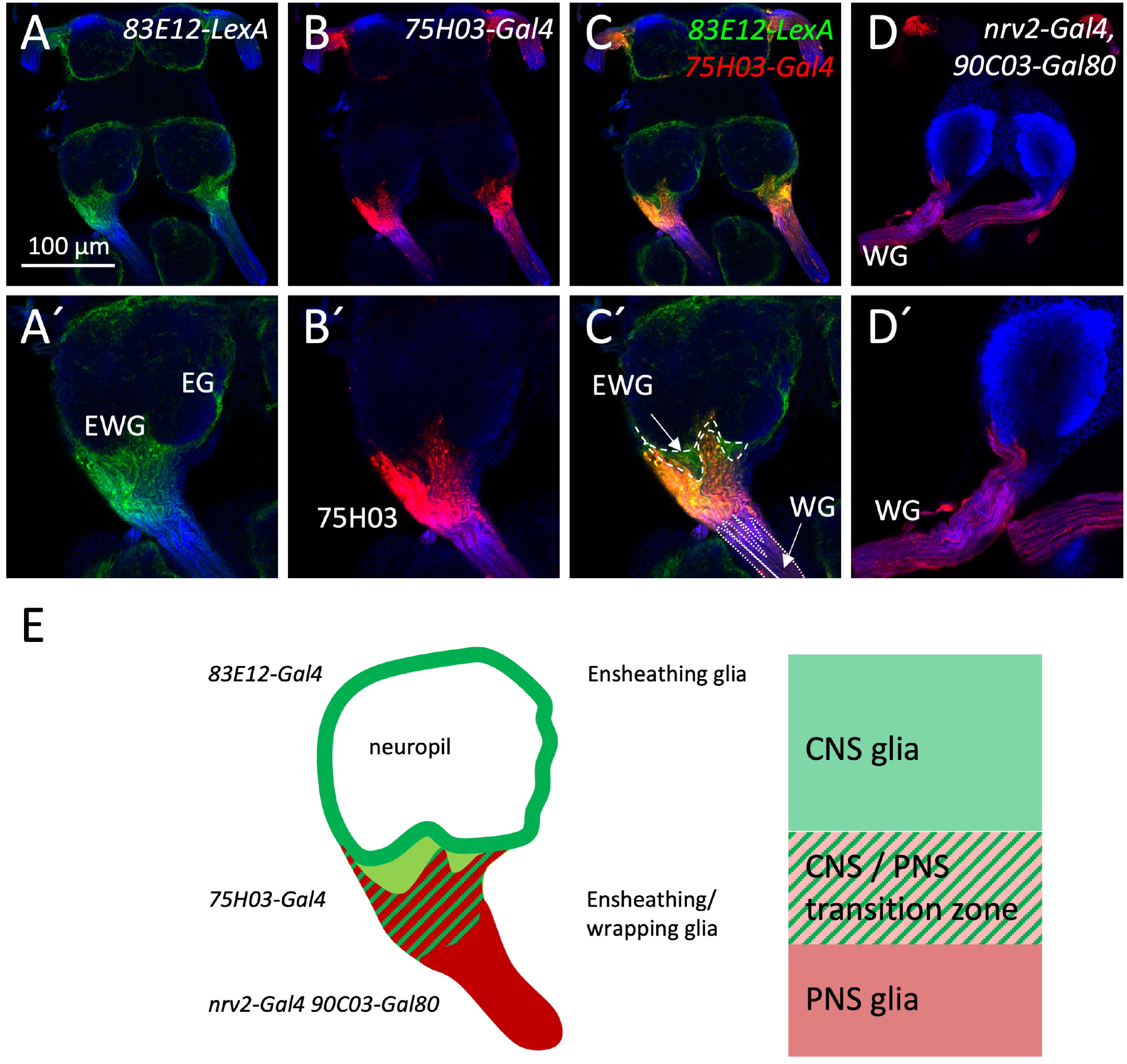
**(A-C)** The tract glial cells as defined by *75H03-Gal4 UAS-GFP* activity, also express the CNS ensheathing glia marker *83E12-LexA LexAop-mCherry*. **(D)** The PNS wrapping glia marker *nrv2-Gal4 90C03-Gal80 UAS-mCherry* labels cells that overlap in their expression domain with the tract glial cells. HRP (blue) labels neuronal membranes. Scale bar is 100 μm. **(E)** Schematic summary of central and peripheral wrapping glial cells in Drosophila. The neuropil is covered by the ensheathing glia. The peripheral axons are wrapped by the peripheral wrapping glia. The *75H03-Gal4* positive glial cells are located in between these two glial cell populations.

**Figure S3.**
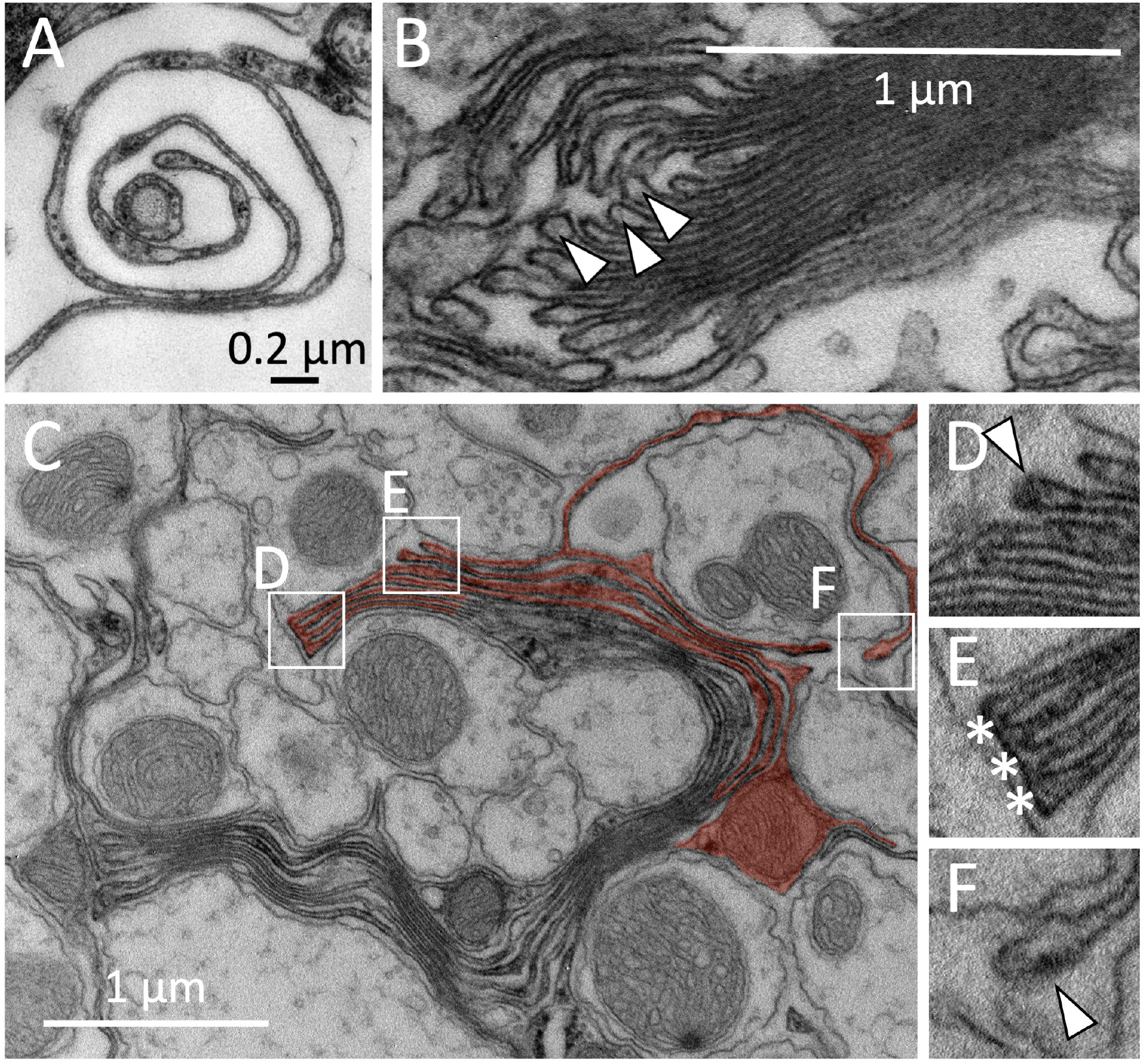
Formation of myelin-like structures in the adult CNS of Drosophila. **(A)** High pressure freezing preparation. Spiral growth of a glial cell process. **(B)** Membrane stack formed by a wrapping glial cell. Note the bulbed growing tips of the glial membrane sheets (arrowheads). **(C-F)** Myelin-like membrane sheets can be connected by comb-like structures. (C) Overview of a multilayered membrane stack around several axons, red shading highlights some of the glial membrane sheets. The arrowhead indicates a bulb structure at the end of the glial membrane sheet (D). In some cases, the ends of the membrane sheets are connected by comb-like structures (asterisks) (E). Growing tip of a wrapping glial cell process that navigated around an axon (ax) (F). Scale bars are as indicated.

**Figure S4.**
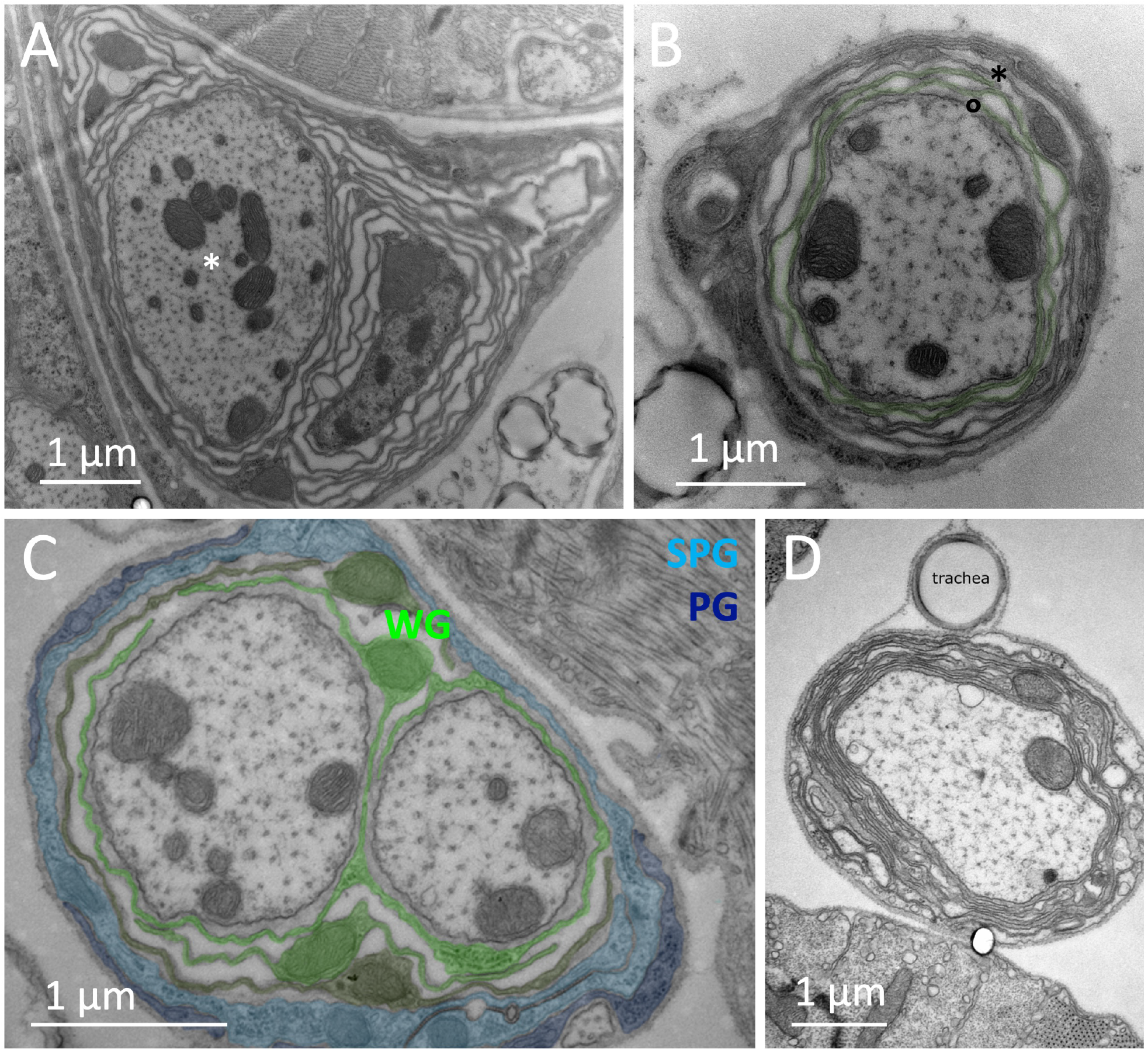
Multilayered myelin-like structures are formed around single axons in the adult nervous system. **(A)** Loosely wrapped glial membranes around one single axon (asterisk). The spacing of the glial membranes resembles the glial lacunae. **(B)** Wrapping around a single axon. The green shaded glial cell process wraps spirally around the central axon. The ends are denoted by the asterisk and the circle. **(C)** Simple wrapping around single axons. The shading indicates the different glial cell types present in the nerve: Wrapping glia WG, perineurial glia PG, subperineurial glia SPG. **(D)** Tight wrapping around a single axon. Unlike the image shown in (A) a close apposition of glial membranes is noted. Scale bars are as indicated.

**Figure S5.**
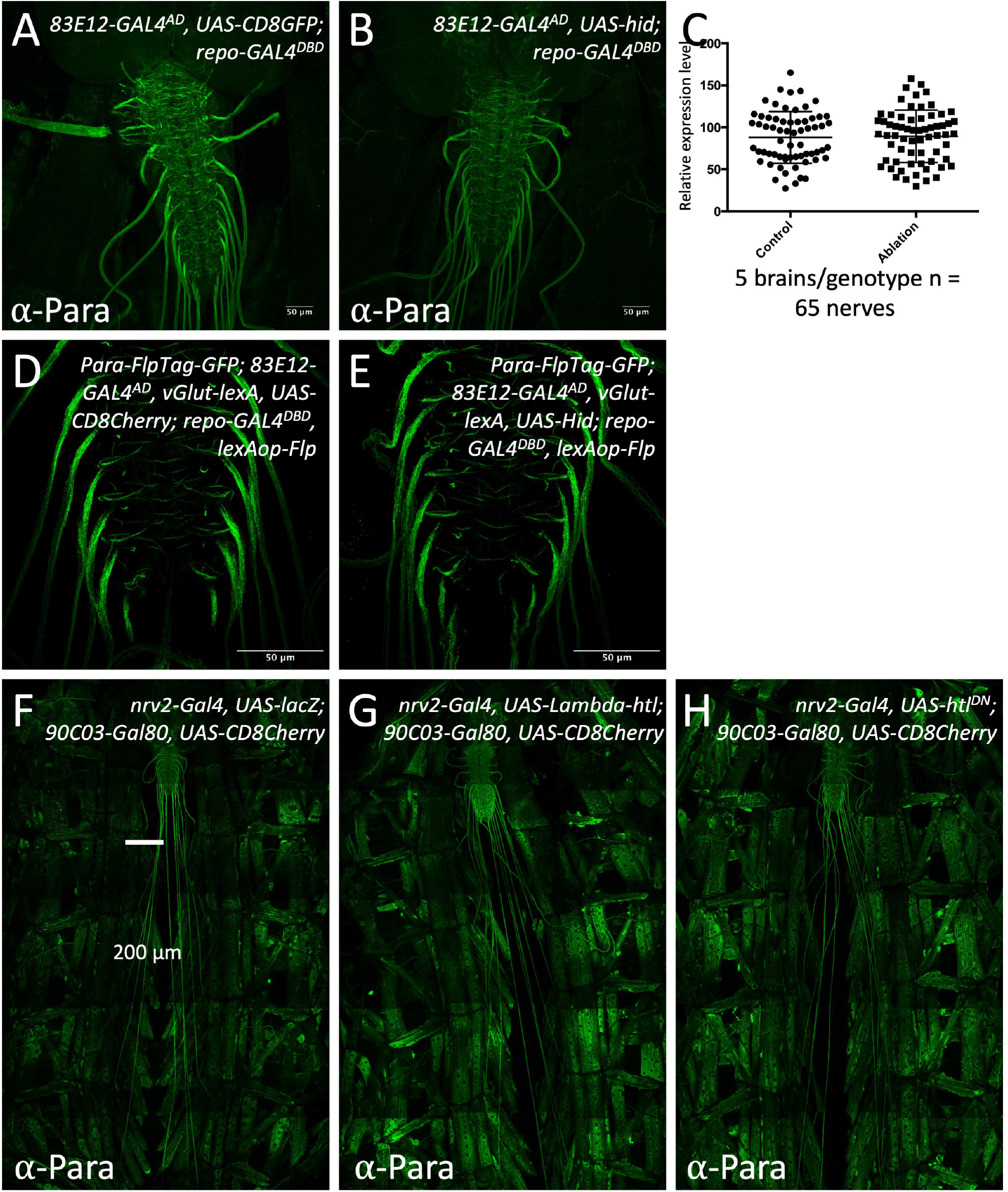
Ablation of central ensheathing glia does not affect positioning of Para at the AIS. CNS preparations of third instar larvae of the genotypes indicated are shown. **(A)** Control larva, expressing CD8GFP under the control of the split Gal4 driver [*83E12-Gal4*^*AD*^, *repo-Gal4*^*DBD*^, *UAS-CD8GFP*] specific for ensheathing glial cells stained for Para protein expression. **(B)** Upon ablation of the ensheathing glia following expression of the proapoptotic gene *hid* no change in the Para expression levels are detected. **(C)** Quantification of the Para staining intensity in control and ensheathing glia ablated larvae. 5 brains per genotypes with 65 nerves were analyzed. **(D)** Control larvae for ensheathing glia ablation using the FlpTag approach. The GFP encoding exon was flipped in all motor neurons using [*vGlut-lexA, lexAop-Flp*]. Note, the pronounced localization of Para^GFP^ at the AIS-like domain of the nerve. **(E)** Upon ablation of the ensheathing glial cells no change in Para localization in motor axons can be detected. **(F-H)** Filet preparations of third instar larvae stained for Para localization. Control larva (F). Upon expression of activated FGF-receptor Heartless no change in Para localization is noted (G). Upon expression of dominant negative Heartless no change in Para localization is noted (H). Scale bars are as indicated.

**Figure S6.**
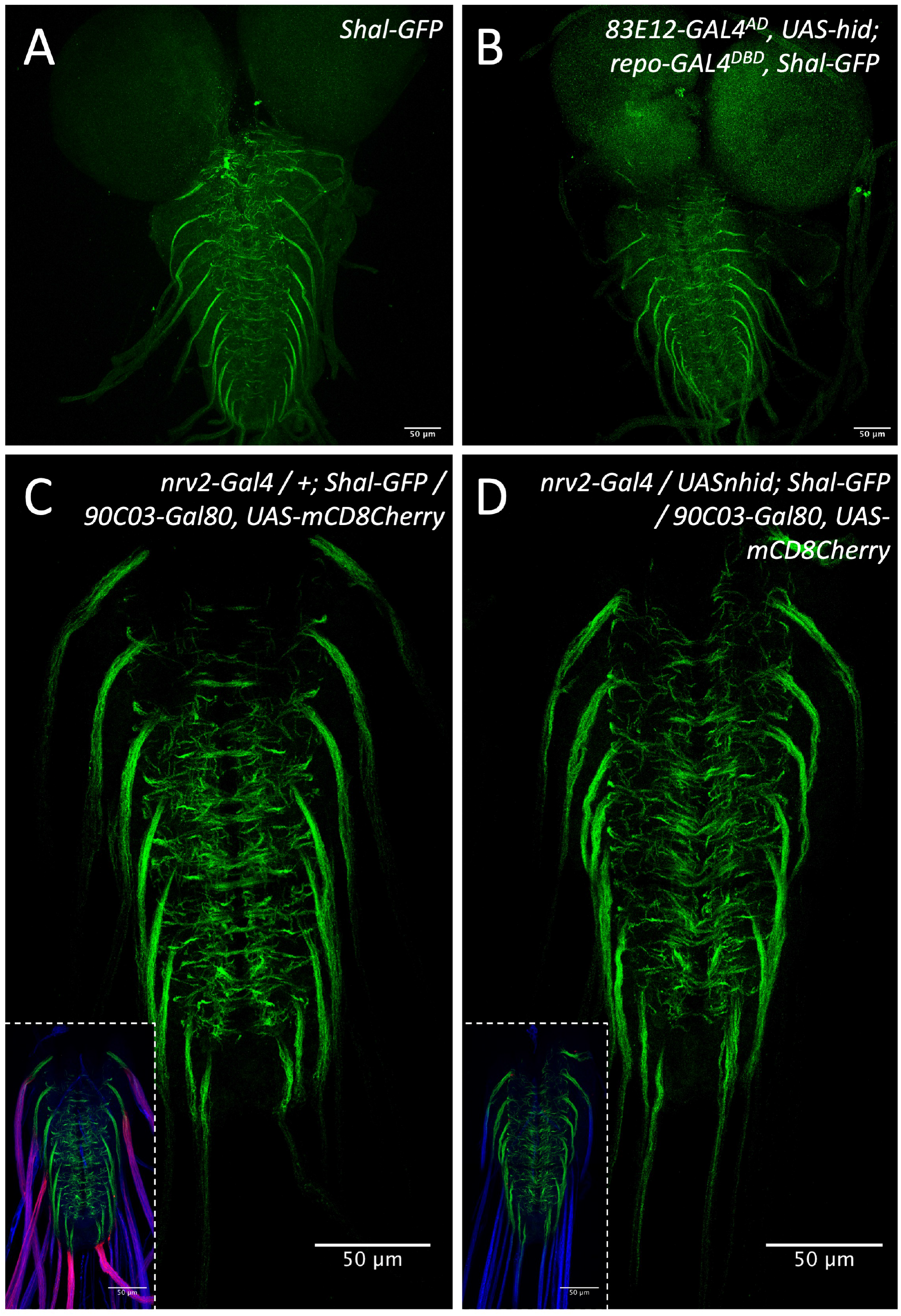
Glia ablation does not affect the localization of the voltage-gated potassium channel Shal. **(A)** Control larva with an endogenously tagged Shal potassium channel. Shal predominantly localizes to the axon initial segment. **(B)** Upon ablation of the ensheathing glia, no change in Shal localization is detected. **(C)** Control larva. The inlay shows co-staining for wrapping glial cell processes (magenta) and HRP to detect neuronal membranes. **(D)** Upon ablation of wrapping glia, no change in Shal localization is detected. Scale bars are as indicated.

**Zip file Sup. Fig.1 with source data**

Four images of western blots, with and without marker bands, are provided.

## References

Allen, M.J., Godenschwege, T.A., Tanouye, M.A., and Phelan, P. (2006). Making an escape: development and function of the Drosophila giant fibre system. Semin Cell Dev Biol 17, 31–41. 10.1016/j.semcdb.2005.11.011.

Amor, V., Feinberg, K., Eshed-Eisenbach, Y., Vainshtein, A., Frechter, S., Grumet, M., Rosenbluth, J., and Peles, E. (2014). Long-term maintenance of Na+ channels at nodes of Ranvier depends on glial contact mediated by gliomedin and NrCAM. Journal of Neuroscience 34, 5089–5098. 10.1523/JNEUROSCI.4752-13.2014.

Anastassiou, C.A., and Koch, C. (2015). Ephaptic coupling to endogenous electric field activity: why bother? Current Opinion in Neurobiology 31, 95–103. 10.1016/j.conb.2014.09.002.

Anastassiou, C.A., Perin, R., Markram, H., and Koch, C. (2011). Ephaptic coupling of cortical neurons. Nature Neuroscience 14, 217–223. 10.1038/nn.2727.

Arancibia-Cárcamo, I.L., Ford, M.C., Cossell, L., Ishida, K., Tohyama, K., and Attwell, D. (2017). Node of Ranvier length as a potential regulator of myelinated axon conduction speed. eLife 6. 10.7554/eLife.23329.

Arvanitaki, A. (1942). Effects evoked in an axon by the activity of a contiguous one. Journal of Neurophysiology 5, 89–108. 10.1152/jn.1942.5.issue-2;requestedJournal:journal:jn;pageGroup:string:Publication.

Bittern, J., Pogodalla, N., Ohm, H., Bruser, L., Kottmeier, R., Schirmeier, S., and Klambt, C. (2021). Neuron-glia interaction in the Drosophila nervous system. Developmental Neurobiology 81, 438–452. 10.1002/dneu.22737.

Castelfranco, A.M., and Hartline, D.K. (2015). The evolution of vertebrate and invertebrate myelin: a theoretical computational study. J Comput Neurosci 38, 521–538. 10.1007/s10827-015-0552-x.

Chandra, P., and Singh, Y.N. (1983). Role of the perineurium and glial lacunar system in nutrition and storage of nutrients in the central nervous system of Spodoptera litura Fabr. (Lepidoptera: Noctulidae) at the time of reorganisation of the neural lamella during metamorphosis. Folia Histochem Cytochem (Krakow) 21, 59–65.

Coggeshall, R.E., and Fawcett, D.W. (1964). The fine structure of the central nervous system of the leech, Hirudo medicinalis. Journal of Neurophysiology 27, 229–289.

Cohen, C.C.H., Popovic, M.A., Klooster, J., Weil, M.-T., Möbius, W., Nave, K.-A., and Kole, M.H.P. (2019). Saltatory Conduction along Myelinated Axons Involves a Periaxonal Nanocircuit. Cell. 10.1016/j.cell.2019.11.039.

Davis, A.D., Weatherby, T.M., Hartline, D.K., and Lenz, P.H. (1999). Myelin-like sheaths in copepod axons. Nature 398, 571. 10.1038/19212.

Dubessy, A.L., Mazuir, E., Rappeneau, Q., Ou, S., Abi Ghanem, C., Piquand, K., Aigrot, M.S., Thétiot, M., Desmazières, A., Chan, E., et al. (2019). Role of a Contactin multi-molecular complex secreted by oligodendrocytes in nodal protein clustering in the CNS. Glia 67, 2248–2263. 10.1002/glia.23681.

Dutta, D.J., Woo, D.H., Lee, P.R., Pajevic, S., Bukalo, O., Huffman, W.C., Wake, H., Basser, P.J., SheikhBahaei, S., Lazarevic, V., et al. (2018). Regulation of myelin structure and conduction velocity by perinodal astrocytes. Proceedings of the National Academy of Sciences 115, 11832–11837. 10.1073/pnas.1811013115.

Eshed-Eisenbach, Y., and Peles, E. (2019). The clustering of voltage-gated sodium channels in various excitable membranes. Developmental Neurobiology. 10.1002/dneu.22728.

Fendl, S., Vieira, R.M., and Borst, A. (2020). Conditional protein tagging methods reveal highly specific subcellular distribution of ion channels in motion-sensing neurons. eLife 9. 10.7554/eLife.62953.

Fontenas, L., and Kucenas, S. (2017). Livin’ On The Edge: glia shape nervous system transition zones. Curr Opin Neurobiol 47, 44–51. 10.1016/j.conb.2017.09.008.

Freeman, S.A., Desmazières, A., Fricker, D., Lubetzki, C., and Sol-Foulon, N. (2016). Mechanisms of sodium channel clustering and its influence on axonal impulse conduction. Cellular and molecular life sciences : CMLS 73, 723–735. 10.1007/s00018-015-2081-1.

Freeman, S.A., Desmazières, A., Simonnet, J., Gatta, M., Pfeiffer, F., Aigrot, M.S., Rappeneau, Q., Guerreiro, S., Michel, P.P., Yanagawa, Y., et al. (2015). Acceleration of conduction velocity linked to clustering of nodal components precedes myelination. Proceedings of the National Academy of Sciences 112, E321–328. 10.1073/pnas.1419099112.

Gisselbrecht, S., Skeath, J.B., Doe, C.Q., and Michelson, A.M. (1996). heartless encodes a fibroblast growth factor receptor (DFR1/DFGF-R2) involved in the directional migration of early mesodermal cells in the Drosophila embryo. Genes & Development 10, 3003–3017.

Günay, C., Sieling, F.H., Dharmar, L., Lin, W.H., Wolfram, V., Marley, R., Baines, R.A., and Prinz, A.A. (2015). Distal spike initiation zone location estimation by morphological simulation of ionic current filtering demonstrated in a novel model of an identified Drosophila motor neuron. PLoS Comput Biol 11, e1004189. 10.1371/journal.pcbi.1004189.

Günther, J. (1976). Impulse conduction in the myelinated giant fibers of the earthworm. Structure and function of the dorsal nodes in the median giant fiber. The Journal of Comparative Neurology 168, 505–531. 10.1002/cne.901680405.

Hama, K. (1959). Some observations on the fine structure of the giant nerve fibers of the earthworm, Eisenia foetida. The Journal of Biophysical and Biochemical Cytology 6, 61–66. 10.1083/jcb.6.1.61.

Hama, K. (1966). The fine structure of the Schwann cell sheath of the nerve fiber in the shrimp (Penaeus japonicus). J Cell Biol 31, 624–632. 10.1083/jcb.31.3.624.

Han, K.S., Guo, C., Chen, C.H., Witter, L., Osorno, T., and Regehr, W.G. (2018). Ephaptic Coupling Promotes Synchronous Firing of Cerebellar Purkinje Cells. Neuron 100, 564-578.e563. 10.1016/j.neuron.2018.09.018.

Hartline, D.K., and Colman, D.R. (2007). Rapid conduction and the evolution of giant axons and myelinated fibers. Current biology 17, R29–35. 10.1016/j.cub.2006.11.042.

Hess, A. (1958). The fine structure and morphological organization of the peripheral nerve-fibres and trunke of the cockroach (Periplaneta americana). Quarterly Journal of Microscopical Science 99, 333–340. 10.1152/jn.1964.27.2.229.

Heuser, J.E., and Doggenweiler, C.F. (1966). The fine structural organization of nerve fibers, sheaths, and glial cells in the prawn, Palaemonetes vulgaris. J Cell Biol 30, 381–403. 10.1083/jcb.30.2.381.

Hodgkin, A.L., and Huxley, A.F. (1952). A quantitative description of membrane current and its application to conduction and excitation in nerve. The Journal of physiology 117, 500–544. 10.1016/j.devcel.2018.10.002.

Igaki, T., Kanuka, H., Inohara, N., Sawamoto, K., Núñez, G., Okano, H., and Miura, M. (2000). Drob-1, a Drosophila member of the Bcl-2/CED-9 family that promotes cell death. Proc Natl Acad Sci U S A 97, 662–667. 10.1073/pnas.97.2.662.

Jegla, T., Nguyen, M.M., Feng, C., Goetschius, D.J., Luna, E., van Rossum, D.B., Kamel, B., Pisupati, A., Milner, E.S., and Rolls, M.M. (2016). Bilaterian Giant Ankyrins Have a Common Evolutionary Origin and Play a Conserved Role in Patterning the Axon Initial Segment. PLoS Genetics 12, e1006457. 10.1371/journal.pgen.1006457.

Kottmeier, R., Bittern, J., Schoofs, A., Scheiwe, F., Matzat, T., Pankratz, M., and Klämbt, C. (2020). Wrapping glia regulates neuronal signaling speed and precision in the peripheral nervous system of Drosophila. Nature communications 11, 4491–4417. 10.1038/s41467-020-18291-1.

Kremer, M.C., Jung, C., Batelli, S., Rubin, G.M., and Gaul, U. (2017). The glia of the adult Drosophila nervous system. Glia 65, 606–638. 10.1002/glia.23115.

Krnjevic, K. (1986). Ephaptic Interactions: A Significant mode of Communications in the Brain. Physiology (Bethesda, Md.) 1, 28–29. 10.1152/physiologyonline.1986.1.1.28.

Kroll, J.R., Saras, A., and Tanouye, M.A. (2015). Drosophila sodium channel mutations: Contributions to seizure-susceptibility. Experimental neurology 274, 80–87. 10.1016/j.expneurol.2015.06.018.

Kucenas, S., Takada, N., Park, H.-C., Woodruff, E., Broadie, K., and Appel, B. (2008). CNS-derived glia ensheath peripheral nerves and mediate motor root development. Nature Neuroscience 11, 143–151. 10.1038/nn2025.

Kucenas, S., Wang, W.D., Knapik, E.W., and Appel, B. (2009). A selective glial barrier at motor axon exit points prevents oligodendrocyte migration from the spinal cord. J Neurosci 29, 15187–15194. 10.1523/jneurosci.4193-09.2009.

Lam, S.S., Martell, J.D., Kamer, K.J., Deerinck, T.J., Ellisman, M.H., Mootha, V.K., and Ting, A.Y. (2015). Directed evolution of APEX2 for electron microscopy and proximity labeling. Nature Methods 12, 51–54. 10.1038/nmeth.3179.

Leech, C.A., and Swales, L.S. (1987). Enzyme effects on the connective tissues of an insect central nervous system. Tissue Cell 19, 587–598. 10.1016/0040-8166(87)90050-4.

Levi, J.U., Cowden, R.R., and Collins, G.H. (1966). The microscopic anatomy and ultrastructure of the nervous system in the earthworm (Lumbricus sp.) with emphasis on the relationship between glial cells and neurons. J Comp Neurol 127, 489–510. 10.1002/cne.901270405.

Maddrell, S.H., and Treherne, J.E. (1967). The ultrastructure of the perineurium in two insect species, Carausius morosus and Periplaneta americana. Journal of Cell Science 2, 119–128.

Mahr, A., and Aberle, H. (2006). The expression pattern of the Drosophila vesicular glutamate transporter: a marker protein for motor neurons and glutamatergic centers in the brain. Gene expression patterns : GEP 6, 299–309. 10.1016/j.modgep.2005.07.006.

Matzat, T., Sieglitz, F., Kottmeier, R., Babatz, F., Engelen, D., and Klämbt, C. (2015). Axonal wrapping in the Drosophila PNS is controlled by glia-derived neuregulin homolog Vein. Development 142, 1336–1345. 10.1242/dev.116616.

Moran, Y., Barzilai, M.G., Liebeskind, B.J., and Zakon, H.H. (2015). Evolution of voltage-gated ion channels at the emergence of Metazoa. Journal of Experimental Biology 218, 515–525. 10.1242/jeb.110270.

Nave, K.-A., and Werner, H.B. (2014). Myelination of the nervous system: mechanisms and functions. Annual Review of Cell and Developmental Biology 30, 503–533. 10.1146/annurev-cellbio-100913-013101.

Nave, K.-A., and Werner, H.B. (2021). Ensheathment and Myelination of Axons: Evolution of Glial Functions. Annual Review of Neuroscience 44, 197–219. 10.1146/annurev-neuro-100120-122621.

Nelson, A.D., and Jenkins, P.M. (2017). Axonal Membranes and Their Domains: Assembly and Function of the Axon Initial Segment and Node of Ranvier. Frontiers in Cellular Neuroscience 11, 136. 10.3389/fncel.2017.00136.

Nern, A., Pfeiffer, B.D., and Rubin, G.M. (2015). Optimized tools for multicolor stochastic labeling reveal diverse stereotyped cell arrangements in the fly visual system. Proceedings of the National Academy of Sciences of the United States of America 112, E2967–E2976. 10.1073/pnas.1506763112.

Peco, E., Davla, S., Camp, D., M Stacey, S., Landgraf, M., and van Meyel, D.J. (2016). Drosophila astrocytes cover specific territories of the CNS neuropil and are instructed to differentiate by Prospero, a key effector of Notch. Development 143, 1170–1181. 10.1242/dev.133165.

Pérez-Moreno, J.J., and O’Kane, C.J. (2019). GAL4 Drivers Specific for Type Ib and Type Is Motor Neurons in Drosophila. G3 (Bethesda, Md.) 9, 453–462. 10.1534/g3.118.200809.

Pogodalla, N., Kranenburg, H., Rey, S., Rodrigues, S., Cardona, A., and Klämbt, C. (2021). Drosophila ßHeavy-Spectrin is required in polarized ensheathing glia that form a diffusion-barrier around the neuropil. Nature communications 12, 6357–6318. 10.1038/s41467-021-26462-x.

Prokop, A., Küppers-Munther, B., and Sánchez-Soriano, N. (2012). Using Primary Neuron Cultures of Drosophila to Analyze Neuronal Circuit Formation and Function. In The Making and Un-Making of Neuronal Circuits in Drosophila, B.A. Hassan, ed. (Humana Press), pp. 225–247. 10.1007/978-1-61779-830-6_10.

R Caré, B., Emeriau, P.-E., Cortini, R., Victor, J.-M., and France;, S.U.U.U.P.U.L.F.-P. (2015). Chromatin epigenomic domain folding: size matters. AIMS Biophysics 2, 517–530. 10.3934/biophy.2015.4.517.

Rasminsky, M. (1980). Ephaptic transmission between single nerve fibres in the spinal nerve roots of dystrophic mice. The Journal of Physiology 305, 151–169. 10.1113/jphysiol.1980.sp013356.

Ravenscroft, T.A., Janssens, J., Lee, P.-T., Tepe, B., Marcogliese, P.C., Makhzami, S., Holmes, T.C., Aerts, S., and Bellen, H.J. (2020). Drosophila Voltage-Gated Sodium Channels Are Only Expressed in Active Neurons and Are Localized to Distal Axonal Initial Segment-like Domains. Journal of Neuroscience 40, 7999–8024. 10.1523/JNEUROSCI.0142-20.2020.

Rey, S., Zalc, B., and Klämbt, C. (2020). Evolution of glial wrapping: A new hypothesis. Developmental Neurobiology. 10.1002/dneu.22739.

Roots, B.I. (2008). The phylogeny of invertebrates and the evolution of myelin. Neuron glia biology 4, 101–109. 10.1017/S1740925X0900012X.

Roots, B.I., and Lane, N.J. (1983). Myelinating glia of earthworm giant axons: thermally induced intramembranous changes. Tissue & Cell 15, 695–709.

Salvaterra, P.M., and Kitamoto, T. (2001). Drosophila cholinergic neurons and processes visualized with Gal4/UAS-GFP. Brain research. Gene expression patterns 1, 73–82. 10.1016/S1567-133X(01)00011-4.

Shneider, M.N., and Pekker, M. (2015). Correlation of action potentials in adjacent neurons. - PubMed - NCBI. Physical Biology 12, 066009.

Sosinsky, G.E., Crum, J., Jones, Y.Z., Lanman, J., Smarr, B., Terada, M., Martone, M.E., Deerinck, T.J., Johnson, J.E., and Ellisman, M.H. (2008). The combination of chemical fixation procedures with high pressure freezing and freeze substitution preserves highly labile tissue ultrastructure for electron tomography applications. J Struct Biol 161, 359–371. 10.1016/j.jsb.2007.09.002.

Stork, T., Engelen, D., Krudewig, A., Silies, M., Bainton, R.J., and Klämbt, C. (2008). Organization and function of the blood-brain barrier in Drosophila. Journal of Neuroscience 28, 587–597. 10.1523/JNEUROSCI.4367-07.2008.

Treherne, J.E., and Schofield, P.K. (1981). Mechanisms of ionic homeostasis in the central nervous system of an insect. J Exp Biol 95, 61–73. 10.1242/jeb.95.1.61.

Trunova, S., Baek, B., and Giniger, E. (2011). Cdk5 regulates the size of an axon initial segment-like compartment in mushroom body neurons of the Drosophila central brain. Journal of Neuroscience 31, 10451–10462. 10.1523/JNEUROSCI.0117-11.2011.

Van Harreveld, A., Khattab, F.I., and Steiner, J. (1969). Extracellular space in the central nervous system of the leech, Mooreobdella fervida. Journal of Neurobiology 1, 23–40. 10.1002/neu.480010104.

Venken, K.J.T., Schulze, K.L., Haelterman, N.A., Pan, H., He, Y., Evans-Holm, M., Carlson, J.W., Levis, R.W., Spradling, A.C., Hoskins, R.A., and Bellen, H.J. (2011). MiMIC: a highly versatile transposon insertion resource for engineering Drosophila melanogaster genes. Nature Methods 8, 737–743. 10.1038/nmeth.1662.

Wigglesworth, V.B. (1959). The Histology of the Nervous System of an Insect, Rhodnius prolixus. Quarterly Journal of Microscopical Science 100, 299–313.

Wigglesworth, V.B. (1960). The Nutrition of the Central Nervous System in the Cockroach Periplaneta Americana L. The Role of Perineurium and Glial Cells in the Mobilization of Reserves. Journal of Experimental Biology 37, 500–512. 10.1242/jeb.37.3.500.

Wilson, C.H., and Hartline, D.K. (2011a). The novel organization and development of copepod myelin. I. Ontogeny. The Journal of Comparative Neurology 519, 3259–3280. 10.1002/cne.22695.

Wilson, C.H., and Hartline, D.K. (2011b). Novel organization and development of copepod myelin. ii. nonglial origin. The Journal of Comparative Neurology 519, 3281–3305. 10.1002/cne.22699.

Yildirim, K., Winkler, B., Pogodalla, N., Mackensen, S., Baldenius, M., Garcia, L., Naffin, E., Rodrigues, S., and Klambt, C. (2022). Redundant functions of the SLC5A transporters Rumpel, Bumpel, and Kumpel in ensheathing glial cells. Biol Open 11. 10.1242/bio.059128.

Yuan, L.L., and Ganetzky, B. (1999). A glial-neuronal signaling pathway revealed by mutations in a neurexin-related protein. Science 283, 1343–1345.

